# VCP increases or decreases tau seeding using specific cofactors

**DOI:** 10.1101/2023.08.30.555637

**Authors:** Sushobhna Batra, Jaime Vaquer-Alicea, Victor A. Manon, Omar M. Kashmer, Andrew Lemoff, Nigel J. Cairns, Charles L. White, Marc I. Diamond

**Affiliations:** Center for Alzheimer’s and Neurodegenerative Diseases Peter O’Donnell Jr. Brain Institute University of Texas Southwestern Medical Center, Dallas, TX; Department of Neurology Peter O’Donnell Jr. Brain Institute University of Texas Southwestern Medical Center, Dallas, TX; Department of Pathology Peter O’Donnell Jr. Brain Institute University of Texas Southwestern Medical Center, Dallas, TX; Department of Biochemistry University of Texas Southwestern Medical Center, Dallas, TX; Department of Clinical and Biological Sciences, Faculty of Health and Life Sciences, University of Exeter, Exeter, United Kingdom

**Keywords:** Tau, APEX2, VCP, p97, Cofactors, Disaggregase, Seeding

## Abstract

**Background:** Neurodegenerative tauopathies may progress based on seeding by pathological tau assemblies, whereby an aggregate is released from one cell, gains entry to an adjacent or connected cell, and serves as a specific template for its own replication in the cytoplasm. *In vitro* seeding reactions typically take days, yet seeding into the complex cytoplasmic milieu can happen within hours. A cellular machinery might regulate this process, but potential players are unknown.

**Methods:** We used proximity labeling to identify factors that control seed amplification. We fused split-APEX2 to the C-terminus of tau repeat domain (RD) to reconstitute peroxidase activity upon seeded intracellular tau aggregation. We identified valosin containing protein (VCP/p97) 5h after seeding. Mutations in VCP underlie two neurodegenerative diseases, multisystem proteinopathy and vacuolar tauopathy, but its mechanistic role is unclear. We utilized tau biosensors, a cellular model for tau aggregation, to study the effects of VCP on tau seeding.

**Results:** VCP knockdown reduced tau seeding. However, distinct chemical inhibitors of VCP and the proteasome had opposing effects on aggregation, but only when given <8h of seed exposure. ML-240 increased seeding efficiency ∼40x, whereas NMS-873 decreased seeding efficiency by 50%, and MG132 increased seeding ∼10x. We screened VCP co-factors in HEK293 biosensor cells by genetic knockout or knockdown. Reduction of ATXN3, NSFL1C, UBE4B, NGLY1, and OTUB1 decreased tau seeding, as did NPLOC4, which also uniquely increased soluble tau levels. Reduction of FAF2 and UBXN6 increased tau seeding.

**Conclusions:** VCP uses distinct cofactors to determine seed replication efficiency, consistent with a dedicated cytoplasmic processing complex that directs seeds towards dissolution vs. amplification.

## Introduction

Neurodegenerative tauopathies include Alzheimer’s and related disorders, and all are caused by intracellular accumulation of pathological tau assemblies (1). In each disorder pathology progresses predictably, at least in part via connected neural networks (2–5). Experimental and observational evidence suggests that this occurs by release of tau aggregates, followed by their entry into a second order cell, a process termed “seeding” which is easily replicated in simple cell models (6–8). The assembly serves as a precise template for its own replication, thereby propagating a specific conformation, or strain (9). This explains the causal linkage of specific tau strains to uniquely induced patterns of pathology in mouse models (10,11), and the diversity of tau filament structures across tauopathies (12).

Amplification of tau assemblies from fibrillar seeds *in vitro* typically takes several days, even under optimized conditions (13,14), whereas in cells this occurs more quickly, sometimes within hours, and in certain cases will faithfully reproduce specific assembly structures (10). Interestingly, no seed amplification assay *in vitro* has yet achieved the fidelity of structural replication that occurs in cells. To enter cells, tau aggregates bind heparan sulfate proteoglycans (HSPGs) on the surface and are taken up via macropinocytosis (15). Most endocytosed tau traffics to the endolysosomal system where it is degraded by lysosomal proteases (16). By contrast, a small fraction of seeding activity steadily enters the cytoplasm with clearance by the proteasome (16). Seeding occurs widely throughout the cytoplasm, and is not necessarily associated with the original aggregates (16). These observations, and others, have led us to speculate that tau seeding is regulatd by an intracellular “machinery” that brings a seed into contact with free tau monomer for amplification. Several proteomics screens from our lab and others have identified proteins associated with established, or chronic, intracellular tau aggregates (17–21), but we still do not understand the factors involved early in the process of seed amplification. In this study, we used proximity labeling at 5h after seed delivery to identify valosin containing protein (VCP/p97) and characterize its regulatory role at the earliest stages of tau aggregation.

## Results

### Proximity labeling of nascent tau aggregates identifies VCP

To identify proteins close to tau as it initiated aggregation, we exploited split-APEX2 (sAPEX2), which renders the enzyme inactive until holoenzyme reconstitution (22). We fused tau repeat domain (RD) containing the disease-associated P301S mutation to APEX2 fragments (AP: aa1-200; EX: aa201-250) each followed by an IRES sequence fused to either blue fluorescent protein (BFP) or mCherry to confirm expression of both constructs. As a negative control, we used tau containing two proline substitutions (I277P / I308P) that prevented formation of beta-sheet structures (23,24). We used tau RD WT as a control against both P301S and dual proline mutants. sAPEX2 alone controlled for background enrichment of any non-tau specific proteins.

We induced tau RD-AP/EX aggregation by Lipofectamine-mediated transduction of cells with full-length (FL), wild-type (WT) tau fibrils. The earliest detectable biotinylation occurred 5h after induction (Supplemental Fig. 1A), so we performed the assay at this time point. Thus, after transduction of cells with tau fibrils, we waited 5h before treating with biotin-phenol (BP) and H_2_O_2_. We then lysed the cells and used streptavidin beads to purify biotinylated proteins. We identified biotinylated proteins using tandem mass-tag mass spectrometry (TMT-MS), pooling data from three independent experiments (Fig. 1A). We identified VCP/p97 as the most significantly enriched hit of this screen (Fig. 1B).

**Figure 1.**
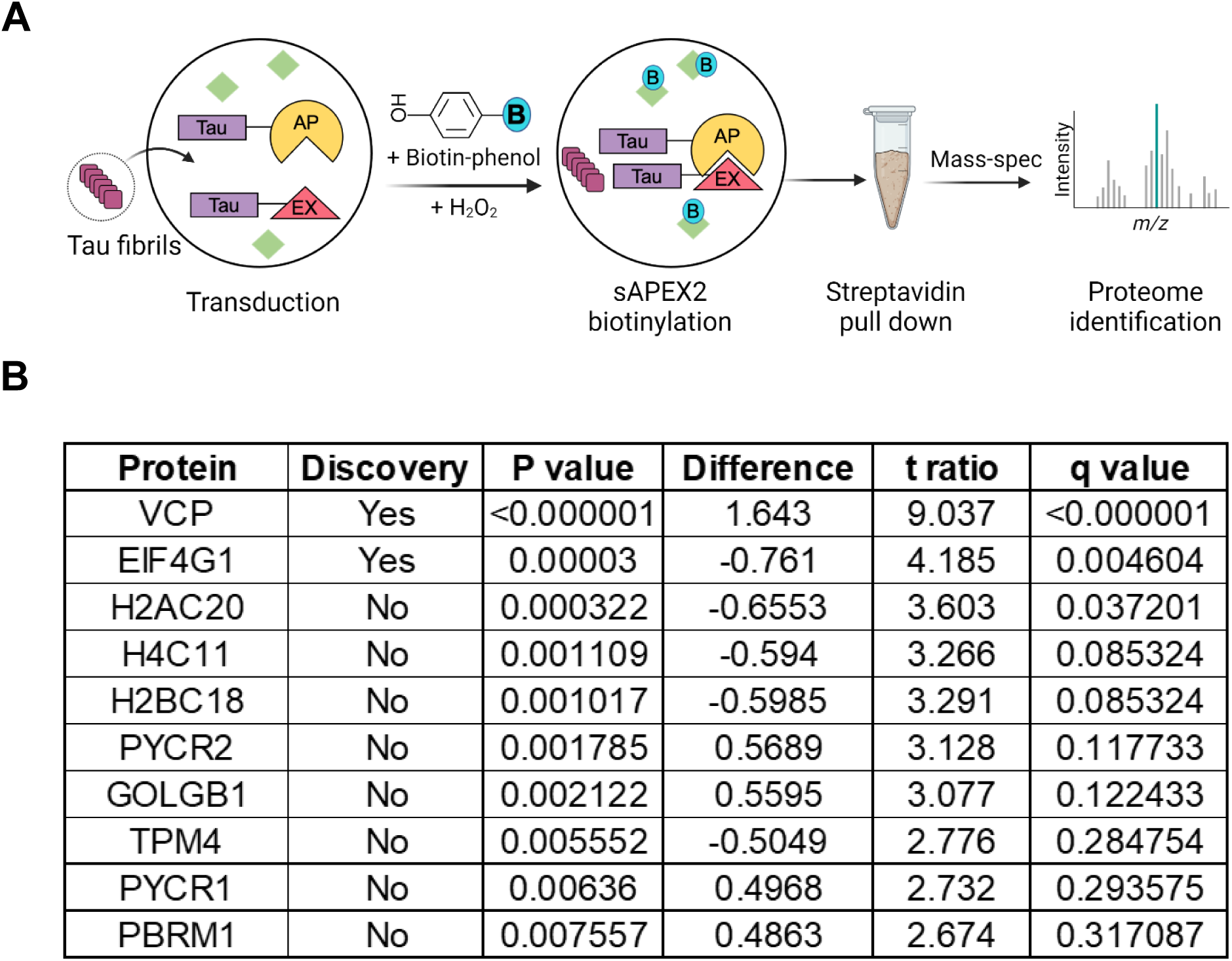
Proximity labeling of nascent tau aggregates identifies VCP. **(A)** Schematic of the TMT-MS study performed for proteomics. **(B)** VCP was identified as the most significant hit enriched in the tau aggregation initiation proteome. Normalized protein abundance ratios for sAPEX2 P301S and sAPEX2 alone (negative control) treated with and without tau fibrils were compared using unpaired t-test on three independent biological replicates; two-stage step-up (Benjamini, Krieger, and Yekutieli), FDR 1.00%). Only top ten proteins of a total of 460 are shown here, based on ascending q values. Difference = enrichment in (sAPEX2 P301S - sAPEX2 alone). Positive difference values indicate enrichment in the aggregation proteome. Statistical significance was determined based on q value.

VCP (known as Cdc48 in yeast and Ter94 in fruit flies) is a AAA+ ATPase with two ATPase domains, D1 and D2. Its N-terminus binds specific cofactor/adaptor proteins that govern its diverse cellular activities (25,26). A dominant mutation in VCP causes vacuolar tauopathy (VT), a neurodegenerative syndrome (27), and other mutations cause multisystem proteinopathy (MSP), with protein aggregation and degeneration in brain, bone, and muscle (28,29). Recently, work from our lab and in an independent collaboration with the Hipp and Hartl laboratories identified VCP associated with insoluble tau aggregates in cells that stably propagate inclusions (20,21). It has been unclear how VCP might regulate intracellular seeding: one group has suggested that it could promote seeding (21), while others have suggested that it prevents seeding (27,30).

### VCP differentially regulates tau seeding

To quantify tau aggregation, we used v2L biosensor cells that overexpress tau RD (P301S) tagged to mClover3 and mCerulean3 (Fig. 2A) (8). We added recombinant tau fibrils to the media in the absence of a transfection reagent to enable HSPG-mediated macropinocytosis (7,15,31) and cytoplasmic seeding (6,32,16), which we quantified using FRET flow cytometry (7). We genetically and pharmacologically modulated VCP activity in the biosensors to test its effect on tau seeding. Knockout (KO) of VCP is lethal, so we first used siRNA-mediated knockdown (KD) (Fig. 2B) and verified it by western blot (Supplemental Fig. 2A). VCP KD reduced tau seeding (Fig. 2C). We also observed increased basal fluorescence of the biosensor cells by microscopy (Supplemental Fig. 2B) and flow cytometry (Supplemental Fig. 2C). This was consistent with VCP-mediated degradation of tau monomer, made the reduction of overall seeding efficiency more remarkable (Fig. 2C, Supplemental Fig. 2D).

**Figure 2.**
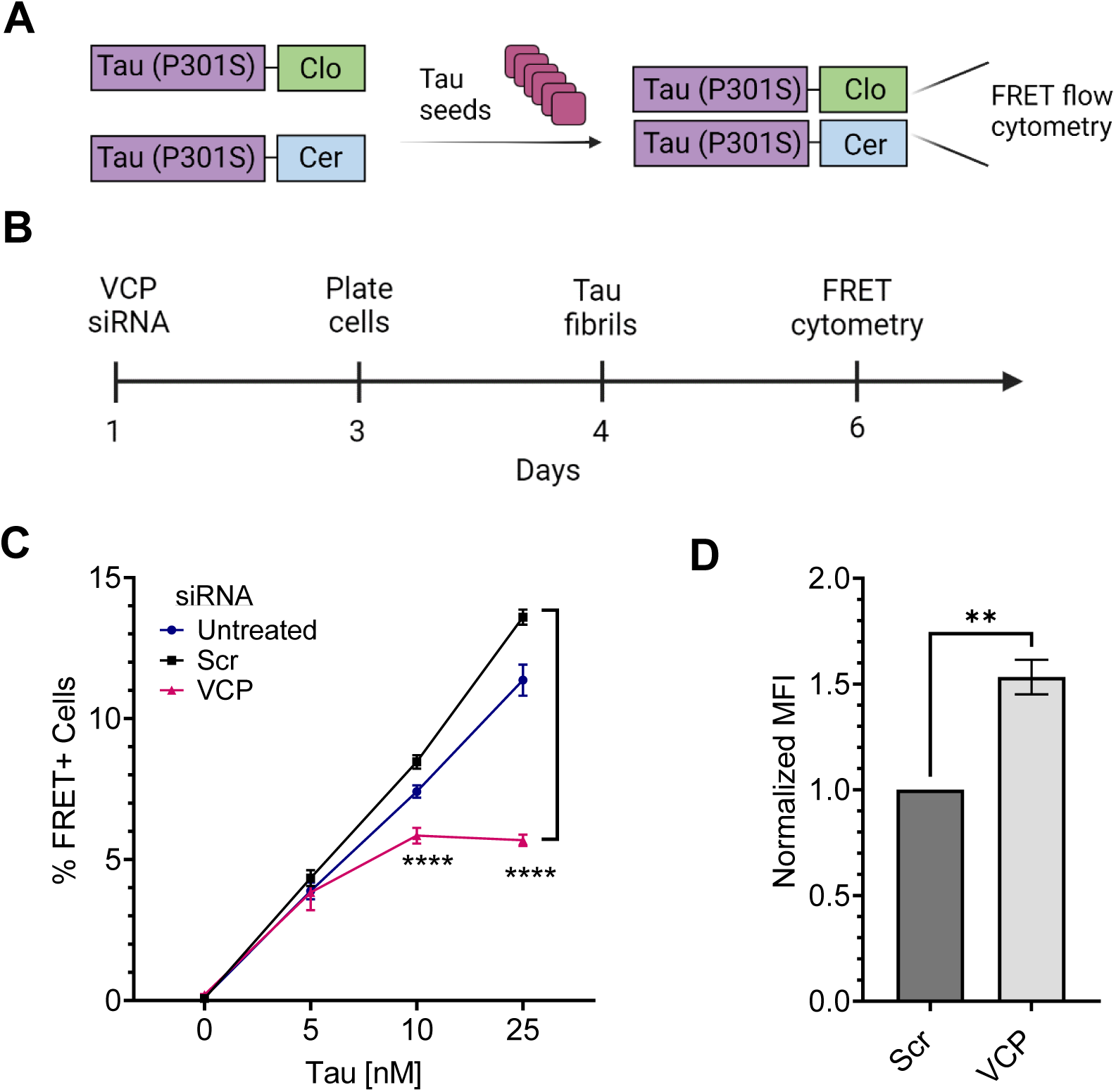
Genetic downregulation of VCP reduces tau seeding. **(A)** Schematic depicting the tau biosensor seeding assay. **(B)** Schematic depicting the siRNA treatment for generating a VCP KD cell line for seeding. **(C)** KD of VCP reduced tau seeding. Error bars represent S.D. Graph is representative of n=3 independent experiments. One-Way ANOVA (Šídák method) with a 95% confidence interval; P value **** < 0.0001 **(D)** VCP KD cells increased uptake of tau fibrils labeled with AF-647, measured by flow cytometry. Error bars represent S.E.M (n=3). Unpaired t-test with a 95% confidence interval; P value ** = 0.003.

To rule out reduction of tau endocytosis, we treated the biosensors with AF-647 (Alexa fluor-647) labeled recombinant tau fibrils for four hours, followed by trypsin digestion to degrade extracellular fibrils. We measured uptake via flow cytometry as per standard protocols (15,31). VCP KD increased tau uptake (Fig. 2D), ruling out diminished endocytosis as the cause of the reduced seeding.

Chronic VCP KD reduced cell proliferation over time and induced toxicity (Supplemental Fig. 2B,C). Thus, we temporarily inhibited VCP using two distinct inhibitors, ML-240 and NMS-873 (Fig. 3A). ML-240 competitively blocks ATP binding at D2, whereas NMS-873 allosterically inhibits VCP by binding to the linker between the D1 and D2 domains (33–35). We pre-exposed the cells to the inhibitors for 1h, then incubated them with tau fibrils for 4h, followed by washout. We then measured induction of seeding after 48h and observed opposing effects. ML-240 increased tau aggregation from approximately 2% to ∼90% (Fig. 3B, E) and speeded aggregation kinetics (Supplemental Fig. 3A). By contrast, NMS-873, reduced tau aggregation (Fig. 3C,E). Because VCP regulates protein degradation via the proteasome (33,36,37), we repeated the study with MG132. This increased tau seeding ∼10x at 48h (Fig. 3D, E), but not as robustly as ML-240 treatment. This agreed with our prior observation that the proteasome mediates cytoplasmic clearance of seeds (16). None of the drugs altered tau uptake (Fig. 3F).

**Figure 3.**
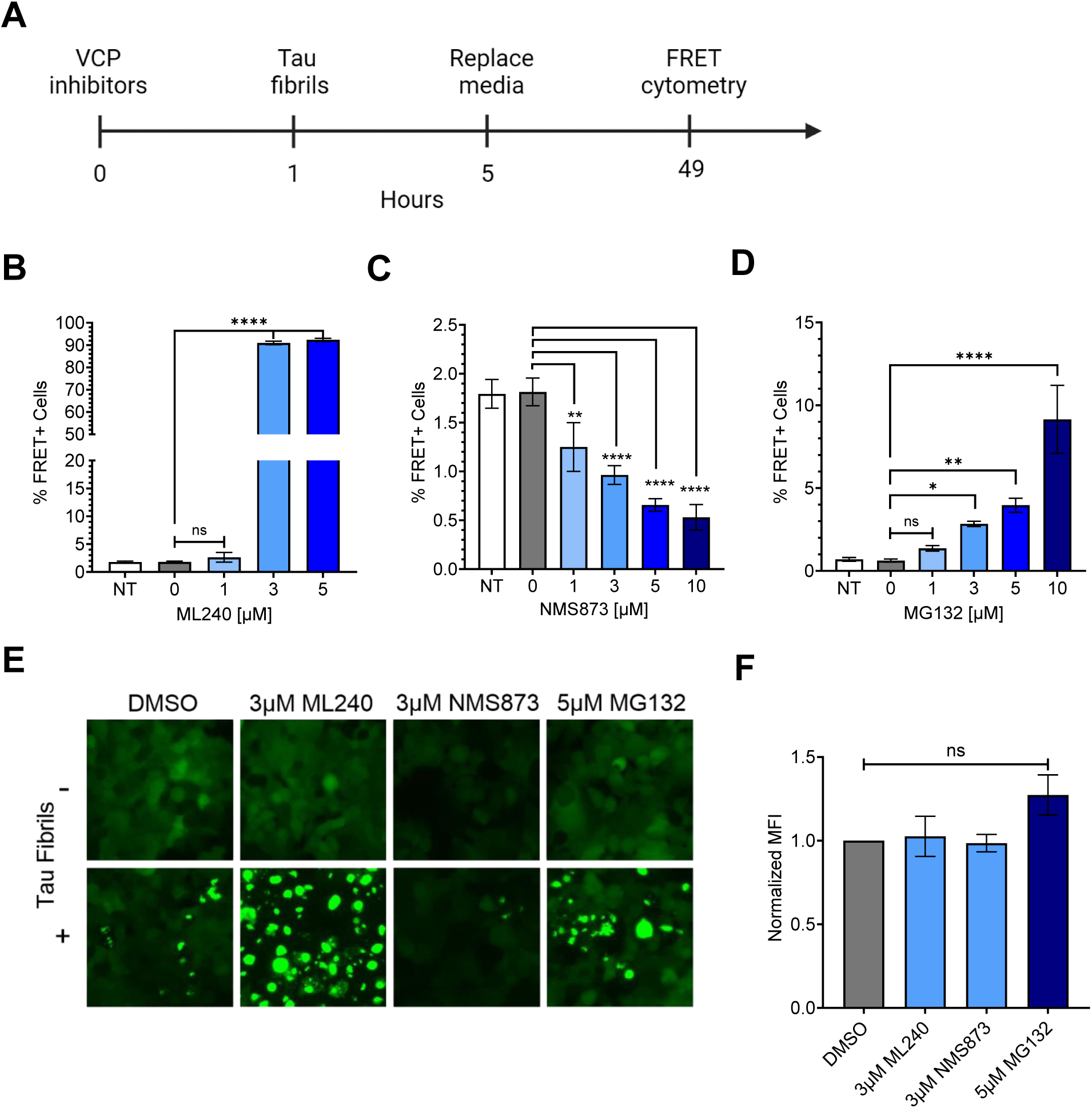
Acute exposure of inhibitors differentially impacts tau aggregation. **(A)** Schematic depicting 1h exposure of tau biosensor cells to inhibitors, followed by 4h of 25nM tau, before washout. **(B)** ML-240 dose-dependently increased tau seeding. P values: ns = 0.44, **** < 0.0001 **(C)** NMS-873 dose-dependently decreased tau seeding. Error bars represent S.D. Representative data of n=3 independent experiments. P values: ** 0.003, **** < 0.0001 **(D)** Proteasome inhibitor MG132 increased tau seeding. P values: ns = 0.85, * =0.04, ** =0.002, **** < 0.0001 **(E)** Fluorescence microscopy confirmed the effects of VCP and proteasome inhibition on tau seeding. **(F)** Drug treatment did not change tau-Alexa 647 uptake as measured by flow cytometry. Error bars represent S.E.M (n=3). P values: ns = 0.996, 0.999, 0.17, in order of bars on the graph. One-Way ANOVA (Šídák method) with a 95% confidence interval.

VCP regulates protein degradation, among other functions, and thus could impact seeding through clearance of aggregates over time. To resolve this issue, we added inhibitors for 7h beginning at different time points after initial seed exposure (Fig. 4A). The inhibitors changed tau seeding only when administered <8h after seed exposure, and we observed no effect on seeding after that timepoint (Fig. 4B, C, D, E; Supplemental Fig. 4A). These results implied that VCP regulates aggregation early in the seeding process.

**Figure 4.**
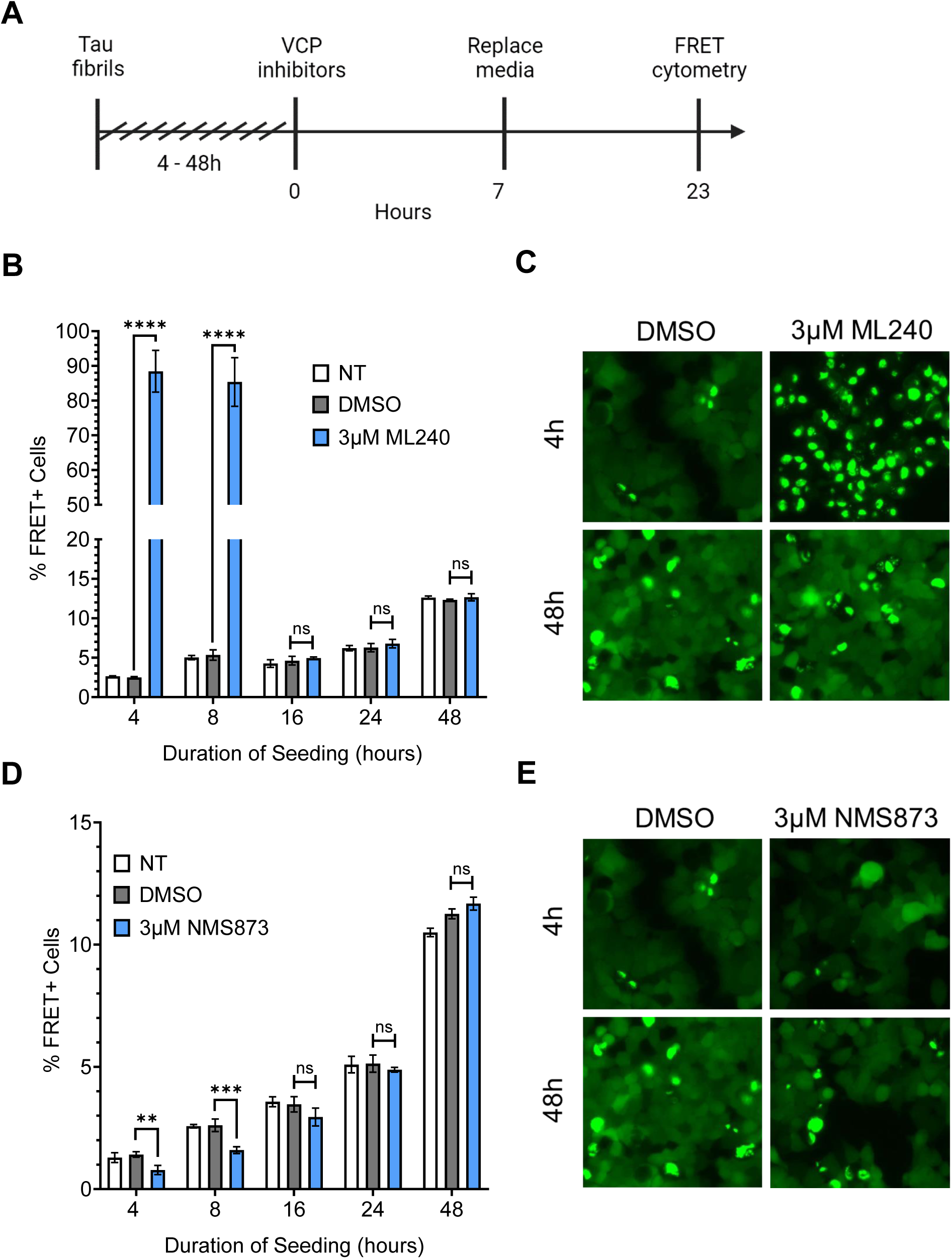
Only early VCP inhibition impacts tau seeding. **(A)** Schematic depicting drug and tau treatments at different time points of the seeding process. **(B)** ML-240 increased tau aggregation ∼16 to 25-fold, but only when administered <8h after seed exposure. P values: **** < 0.0001, ns (16hr) = 0.59, ns (24hr) = 0.43, ns (48hr) = 0.36 **(C)** Representative images (20x magnification). **(D)** NMS-873 decreased tau seeding by ∼50%, but only when administered <8h after seed exposure. P values: ** =0.004, *** =0.0004, ns (16hr) = 0.08, ns (24hr) = 0.87, ns (48hr) = 0.05. **(E)** Representative images (20x magnification). Error bars represent S.D. Representative data of n=3 independent experiments. One-Way ANOVA (Šídák method) with a 95% confidence interval.

### ML-240 increases tauopathy brain lysate seeding

To test the effects of VCP inhibitors on a physiological tau seed source, we tested tauopathy brain lysates from AD (Alzheimer’s disease) and CBD (corticobasal degeneration). We treated biosensor cells with ML-240 and NMS-873 as described above, after which they were exposed to AD and CBD patient brain lysates without a transduction reagent. We used Huntington’s disease brain lysate as a negative control. ML-240 increased AD and CBD seeding by ∼10x (Fig. 5A,C). Since the lysates induced very low seeding in the absence of a transduction reagent, we could not test the effects of NMS-873.

**Figure 5.**
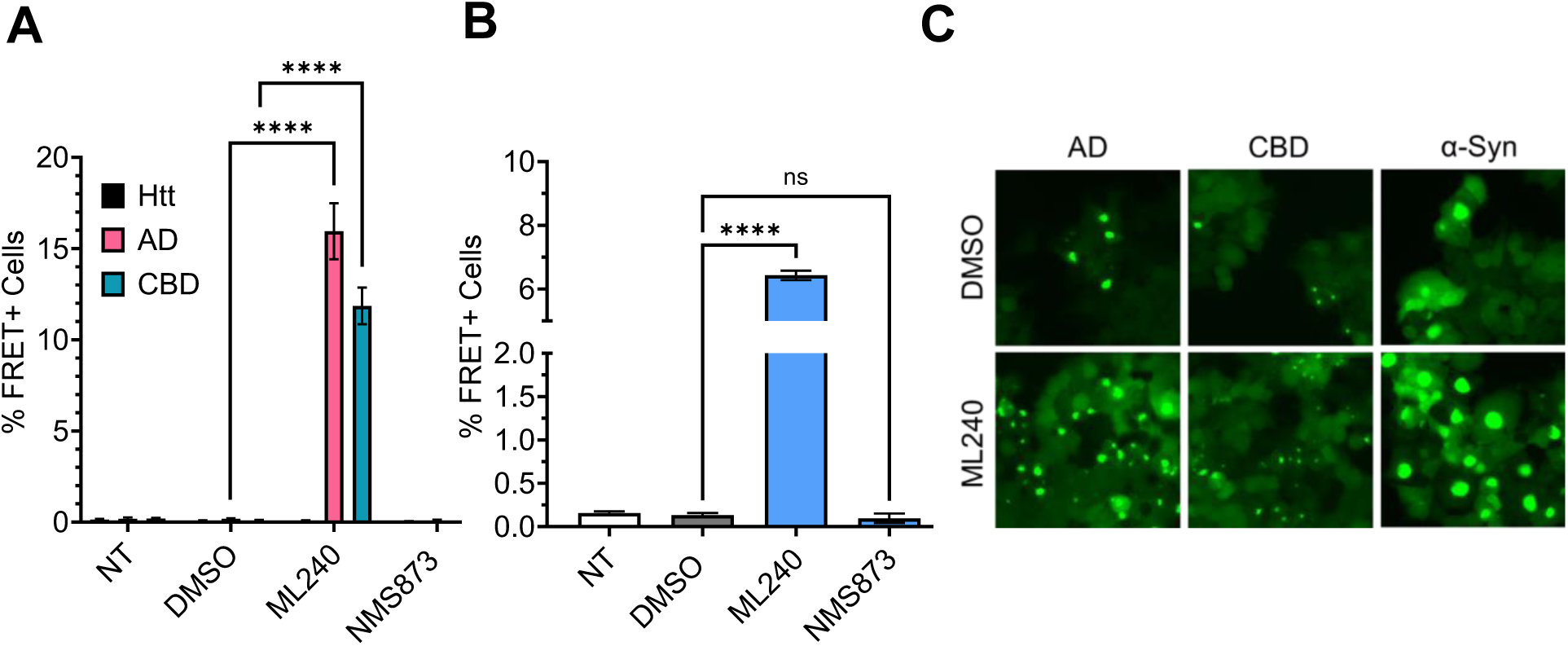
ML-240 enhances seeding by tauopathy brain lysates and recombinant α-synuclein. AD and CBD brain lysates were seeded onto v2L biosensors; recombinant a-synuclein was seeded onto α-synuclein (A53T) biosensors. **(A)** ML-240 increased seeding by AD and CBD brain samples. No seeding from Huntington disease brain lysate (Htt) was observed on tau biosensors. P value, **** < 0.0001 **(B)** ML-240 increased α-synuclein seeding in A53T synuclein biosensors. Error bars represent S.D. Representative data for n=3 independent experiments. P value, **** < 0.0001 **(C)** Representative fluorescence microscopy images for effects of ML-240 on tauopathy and α-synuclein seeding (20x magnification). One-Way ANOVA (Šídák method) with a 95% confidence interval.

### ML-240 increases α−synuclein seeding

To test whether VCP inhibition had similar effects on protein aggregation of another known amyloid, we used biosensors that express FL α-synuclein containing a disease-associated mutation (A53T) fused to cyan and yellow fluorescent proteins (7,38). ML-240 increased the seeding by α-synuclein fibrils ∼6x (Fig. 5B,C). Since the α-synuclein fibrils induced very low seeding in the absence of a transduction reagent, we could not test the effects of NMS-873.

### VCP co-factors regulate tau aggregation

Multiple co-factors generate specificity for VCP’s myriad cellular functions (25,39,40). To identify those which regulated seeding, we individually knocked out or reduced expression of 30 known cofactors in v2L tau biosensor cells (41). These cofactors and their proposed functions have been listed in Table 1.

**Table 1:**
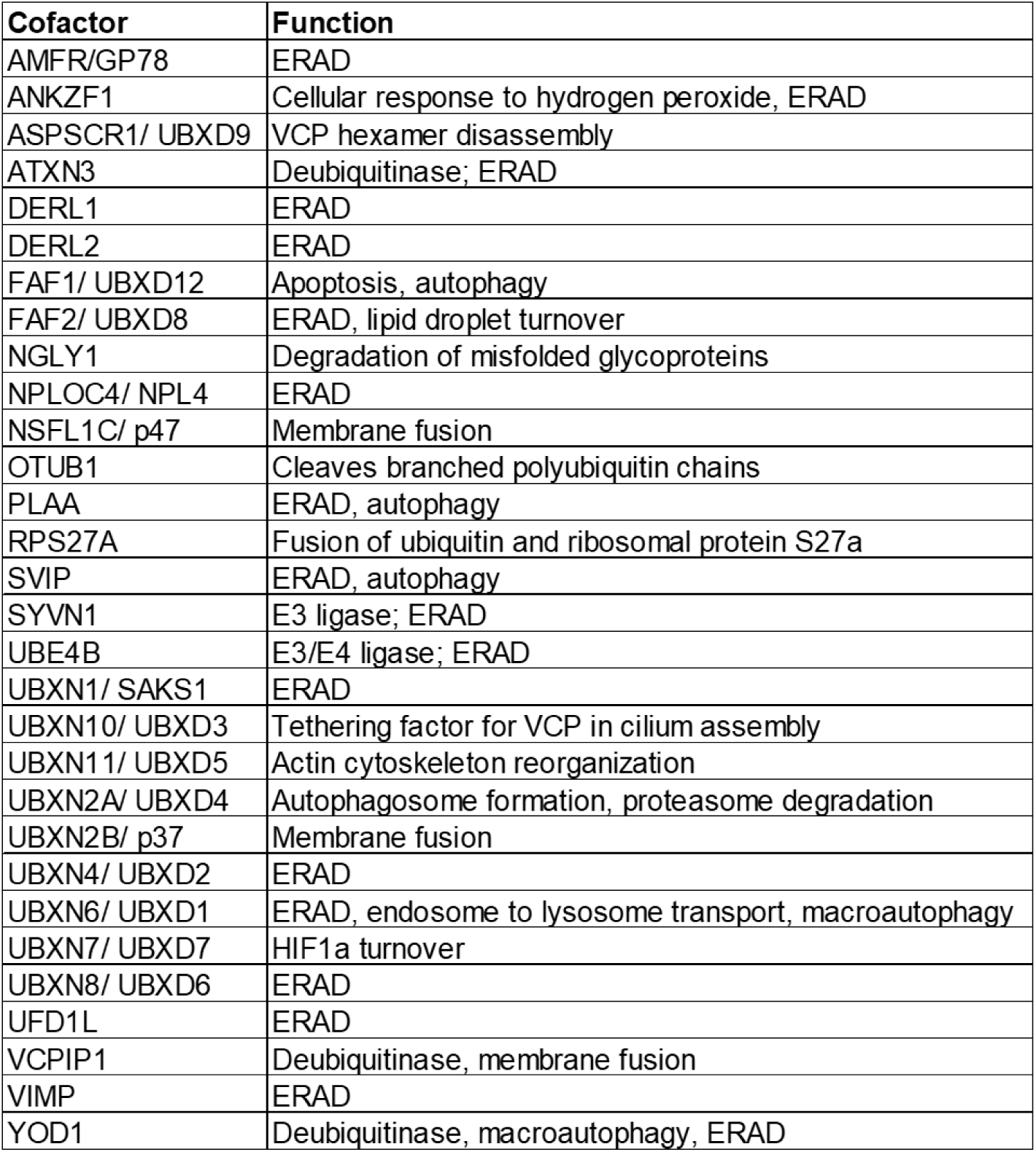
List of VCP cofactors and their proposed functions.

For CRISPR/Cas9 KO, we used four gRNAs per gene from the Brunello library (42), compared to 4 non-targeting guides (NTG) as a negative control. For genes that were toxic upon KO, we used siRNA-mediated KD, with scrambled (Scr) siRNA as a negative control. KO or KD of most cofactors did not change tau seeding (Supplemental Fig. 5A). RPS27A was the only cofactor for which both KD and KO were lethal and thus we could not determine its effects on seeding.

Knockout of UBXN6 increased tau seeding but the effect was most pronounced at higher tau concentrations (Supplemental Fig. 5B). KO of FAF2 alone increased tau seeding (Fig. 6A), and even induced spontaneous aggregation in the biosensors in the absence of any exogenous tau fibrils, based on microscopy (Supplemental Fig. 5C) and FRET flow cytometry (Supplemental Fig. 5D). In contrast, KO of the deubiquitinase ATXN3, and the E3/4 ligase UBE4B, suppressed seeded tau aggregation (Fig. 6B,D). KO of NSFL1C also reduced seeding (Fig. 6C). We validated each effective KO by western blot (Supplemental Fig. 5E). No cofactor KO changed tau uptake (Fig. 6E).

**Figure 6.**
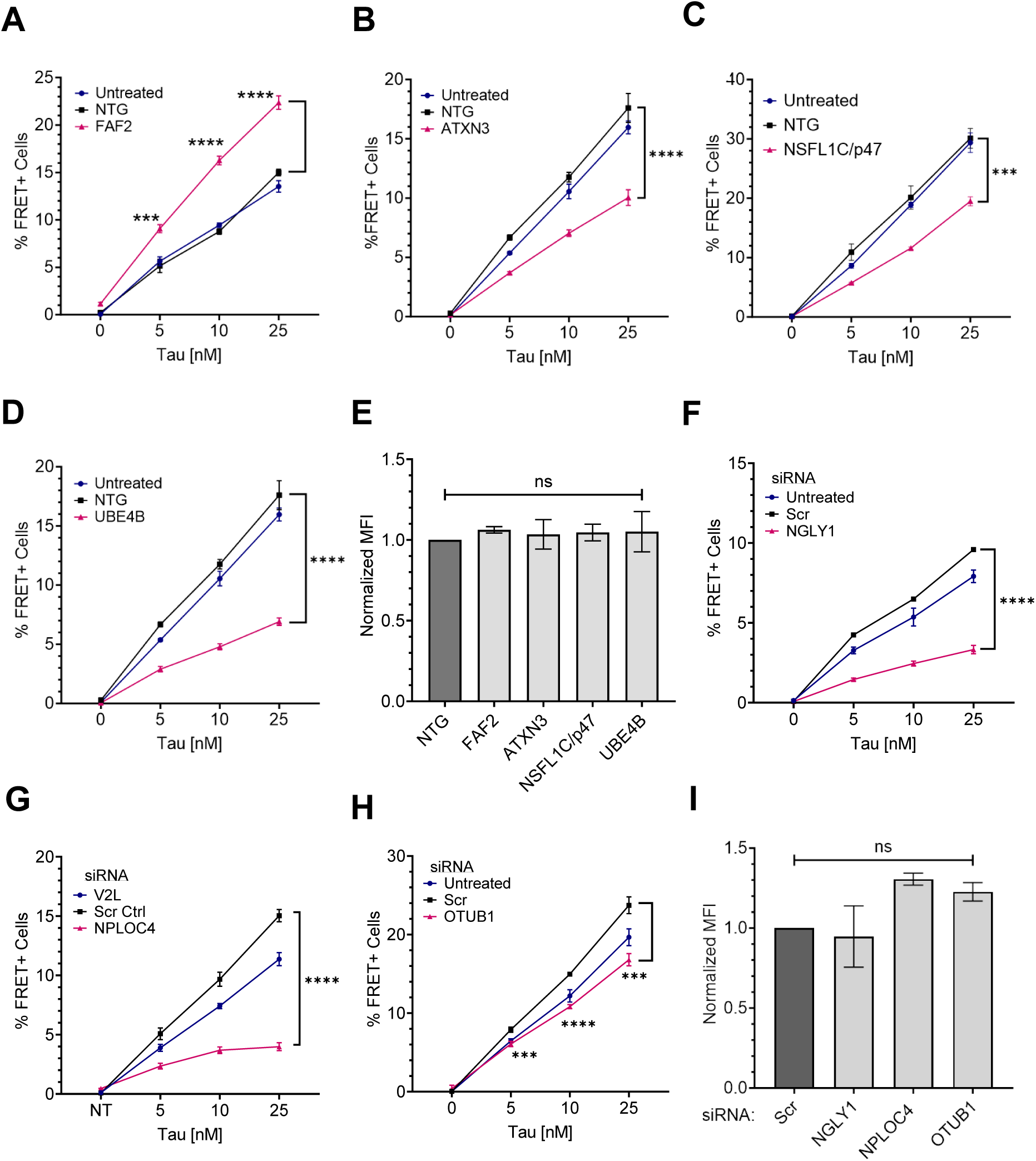
VCP cofactors differentially regulate tau seeding. VCP cofactors were either knocked out via CRISPR/Cas9 (**A-E**) or knocked down via siRNA (**F-I**) in v2L biosensors prior to exposure to increasing amounts of tau fibrils. **(A)** Knockout of FAF2 increased tau seeding whereas knockout of **(B)** ATXN3, **(C)** NSFL1C, and **(D)** UBE4B reduced tau seeding. P values: FAF2 (*** =0.0001, **** < 0.0001); ATXN3 (**** < 0.0001); NSFL1C (*** =0.0002); UBE4B (**** < 0.0001). **(E)** Cofactor KO did not affect tau uptake. P values: ns = 0.98, 0.998, 0.995, 0.99, in order of bars on the graph. **(F)** Knockdown of NGLY1, **(G)** NPLOC4, and **(H)** OTUB1, decreased tau seeding. P values: NGLY1(**** < 0.0001); NPLOC4 (**** < 0.0001); OTUB1 (*** =0.0004, **** < 0.0001, *** =0.0001). **(I)** KD did not affect tau uptake. P values: ns = 0.98, 0.19, 0.39, in order of bars on the graph. Graphs are representative of n= 3 separate experiments. Error bars represent S.D. for (A-D; F-H), S.E.M. for (E,I). One-Way ANOVA (Šídák method) with a 95% confidence interval.

siRNA identified three cofactors whose KD decreased seeding: NGLY1, NPLOC4, and OTUB1 (Fig. 6F,G,H). Remarkably, KD of NPLOC4 increased tau levels in the biosensors (Supplemental Fig. 5F,G) and yet reduced the actual number of aggregates as observed under the microscope (Supplemental Fig. 5F) and as FRET signal on the flow cytometer (Supplemental Fig. 5H). NPLOC4 KD increased inclusion size, whereas NGLY1 KD reduced fluorescence and created smaller puncta (Supplemental Fig. 5F). We validated each effective KD by western blot (Supplemental Fig. 5I). No cofactor KD changed tau uptake (Fig. 6I).

## Discussion

To identify factors that participate in tau seed amplification, we used proximity labeling to identify VCP, which has been genetically and biochemically linked to tau aggregation by other studies (20,21,27), and to inhibition of α-synuclein and TDP-43 seeding (30). Chemical manipulations of VCP up- and down-regulated tau seeding, but only within the first 8h of seed exposure. We observed these effects also with brain-derived tau seeds and recombinant α-synuclein. We identified selected VCP cofactors that participated in this differential regulation, suggesting that a complex within the cell processes incoming seeds, either to decrease or increase their replication efficiency.

Synthesizing our recent data with prior work on VCP and an analogous yeast disaggregase, Hsp104, which controls yeast prion replication (43), we hypothesize that VCP couples two functions in its regulation of seeding: extraction of monomer from the amyloid assembly, and subsequent proteasome-mediated degradation (Fig. 7). If this occurs at the fibril terminus, the seed will not amplify efficiently as there will be no increase in free ends. Conversely, extraction of tau monomer from the middle of the fibril would break it into smaller fragments, increasing the number of free ends. A similar mechanism of yeast fibril fragmentation has been previously described for Hsp104 (44) and also for VCP-mediated tau fibril disaggregation (21). We further propose that the chemical inhibitors and cofactors identified in this study differentially impact these functions either to increase or decrease seeding. This model for VCP makes specific predictions that will require further testing, and reconciles observations by us and others that VCP could prevent (27,30,45) or promote (21) seeding by tau and other amyloid proteins.

**Figure 7.**
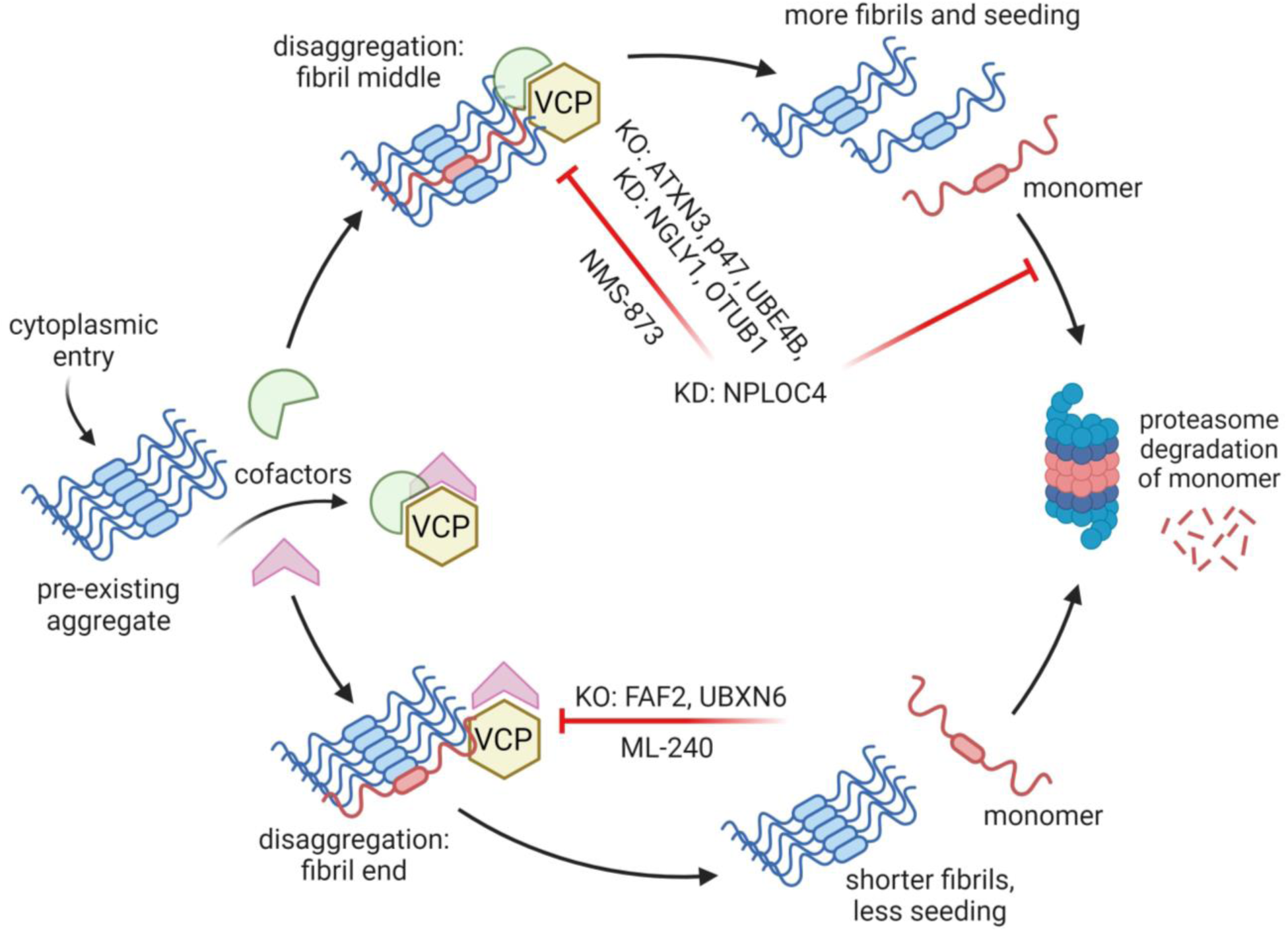
Model of VCP regulation of the fate of tau seeds. VCP acts on tau seeds that enter the cytoplasm, either to promote degradation or amplification. Disaggregase activity of VCP removes monomer for degradation. This can occur at the end of filaments, which would decrease seeding, or from within, which would increase free ends and promote seeding. The effects of NPLOC4, which increases overall tau levels but decreases seeding indicates that these processes are separable. Chemical inhibitors and cofactors bias the process towards differential processing paths. Model based on Saha et al. (21). Image created using biorender.com.

### VCP controls the fate of seeds

Our prior work previously defined a trafficking pathway for tau seeds that delivers them to the cytoplasm, where they are cleared by the proteasome (16). In simple cells such as HEK293T, seeding efficiency is relatively low, consistent with robust clearance. However, we observed a dramatic increase in seeding efficiency for recombinant and patient-derived seeds in the presence of ML-240, a VCP inhibitor that targets the D2 ATPase. Conversely, NMS-873, an allosteric inhibitor, reduced tau seeding by ∼50%, as did knockdown of VCP. These effects were independent of other cellular mechanisms that might be expected to influence seeding: uptake and steady-state tau levels. In fact, we observed a seemingly paradoxical effect in the context of knockdown of VCP and NPLOC4: increased tau steady state with less seeding. We propose that VCP controls the fate of seeds by two distinct activities, each based on extracting a tau monomer from an assembly. If taken from the end of a fibril, this would lead to reduced seeding, as the assemblies would be diminished in size, and would be cleared more efficiently; if taken from within the fibril, this would promote seeding, as there would be fibril cleavage, with more free ends to serve as templates. This model of a balance of disassembly vs. degradation is based on studies of the yeast prion disaggregase, Hsp104, which regulates prion propagation and dissolution (43). We note that others have proposed different models for VCP effects. Darwich et al. suggested that vacuolar tauopathy mutations function by reducing VCP disaggregase activity (27), and hence clearance of aggregates; whereas Zhu et al. concluded that VCP surveillance of permeabilized endosomes might be its primary role (30). Because we have found no evidence that tau aggregates permeabilize endosomes (16), and that different inhibitors can either increase or decrease seeding, we have proposed a distinct model for VCP activity, at least for tau.

### VCP functions early in the seeding process

We designed the proximity labeling to identify proteins adjacent to newly formed tau aggregates at the earliest possible time point, and identified VCP as the single, most reliable hit. We suspect this was because APEX2 holoenzyme reconstitution was limited, restricting the labeling efficiency. Despite the clear identification of VCP with mature aggregates based on our work and that of others (20,21), our treatment/washout studies indicate a critical role of VCP in processing of tau seeds as they first enter the cytoplasm. While there are conflicting reports about how seeds might exit the endolysosomal system (16,32), we have previously found that tau seeds in the cytoplasm are cleared by the proteasome (16). Thus, we propose that VCP contacts seeds that exit endocytic vesicles in the cytoplasm to facilitate their amplification or clearance.

### VCP cofactors dictate the fate of tau seeds

Multiple efforts have attempted unsuccessfully to directly target VCP with small molecules to treat cancer (46). Our observations with ML-240 and NMS-873, which we originally assumed would have the same effect on tau seeding, highlighted the mechanistic complexity of this enzyme. Indeed, we found that specific cofactors were necessary for VCP to control seed amplification vs. destruction. Knockout of FAF2, also known as UBXD8, strongly enhanced tau seeding. It also caused spontaneous aggregation in the biosensors, which we had never before observed. FAF2 was recently reported to facilitate VCP-dependent disaggregation of stress granules (47), and we hypothesize in this case that FAF2 facilitated removal of tau monomer from fibril ends. By contrast, knockdown of NPLOC4 increased tau levels, yet inhibited seeding. This was consistent with prevention of seed amplification, possibly by inhibiting removal of tau from within fibrils for degradation. Genetic deletion of ATXN3 (Ataxin 3), a deubiquitinase, also suppressed tau seeding without affecting tau levels. Ataxin3 is proposed to facilitate ubiquitinated substrate release from VCP by cleaving the ubiquitin chains to a minimum length for proteasomal degradation (48). However, its exact role in processing VCP substrates remains unclear. In the absence of its deubiquitinase, Cdc48 cannot thread its substrate through its disaggregation core (49). Thus, ATXN3 KO might also prevent fragmentation of assemblies, without directly inhibiting monomer degradation. Taken together, our results point to a complex molecular machine, likely with cell-specific components, that determines the fate of tau seeds.

### Multiple roles of VCP in degenerative disorders

Distinct VCP mutations cause MSP with ubiquitinated aggregates of TDP-43 (50,51) or VT (27). MSP mutants have been proposed to be hyperactive (52,53) whereas VT is proposed to result from VCP hypoactivity (27). It is also suggested that MSP mutants induce a conformational change that reduces VCP binding to its cofactors (54,48,53) which might explain the multiple pathways affected in disease. It appears that *in vitro* experiments of ATP hydrolysis in isolation fail to account for the complexity of VCP function in a cellular setting.

Recent work has suggested that VCP regulates protein aggregation of TDP-43, α-synuclein, and tau through multiple potential mechanisms including autophagy, endolysosomal degradation, and disaggregation (27,55,30,21). Our findings add a new dimension by highlighting the role of VCP early in the tau aggregation process, and specific cofactors that differentiate its activities. Indeed, others have proposed UBXN6/UBXD1 as a VCP cofactor that regulates α-synuclein seeding in neurons (30). It is possible that neurons might differentially utilize these or potentially different cofactors to regulate tau seeding, and this will await additional comprehensive study. It seems likely that modulation of specific sub-functions of VCP by inhibiting cofactor interactions might be the most productive approach to therapeutic targeting of an enzyme that is otherwise critical for cell viability.

## Conclusion

This study highlights a critical role of VCP in dictating the fate of tau seeds for either amplification or degradation early in the seeding process. We have identified for the first time VCP cofactors that can specifically regulate tau seeding in a cellular model. Our findings implicate VCP as a master regulator of mammalian amyloids in degenerative disorders and provide an avenue for developing novel and highly specific anti-tau therapeutics.

## Methods

### Cell Culture

HEK293T cells were obtained from ATCC and used to make all cell lines. Cells were maintained in Dulbecco’s DMEM with 10% fetal bovine serum, 1% penicillin-streptomycin, and 1% GlutaMax. v2L tau biosensor cells were used for all seeding assays. Details on these biosensors have been recently published (8). Cell lines were frequently checked for mycoplasma contamination (Venor-GEM Mycoplasma Detection kit).

### Proteomics Screen

T225 flasks were coated with 10mL of 1x poly-D-lysine (PDL) for 2-3h in the incubator and rinsed with PBS before plating cells. 22 million cells were plated in 25ml/T225 flask and allowed to settle overnight. The following day, cells were treated with 50nM tau + Lipofectamine-2000 complexes (or 50nM α-synuclein + Lipofectamine as a negative control) which were incubated for 20min at RT prior to addition to cells. Cells were incubated with the fibrils for 5h. Thirty minutes before the 5h time point, cells were treated with BP (biotin phenol) at a final concentration of 500µM at 37°C. At 5h, cells were treated with H_2_O_2_ at a final concentration of 1mM and the flasks were agitated at RT for 1min. The biotinylation reaction was quenched with the quenching buffer followed by three additional rinses with it. Quenching buffer was also used to scrape the cells to collect the cell pellets. This buffer was prepared as previously described in the APEX2 labeling protocol (56).

### On-Bead Trypsin Digestion

Protein concentrations were normalized across all the samples (∼1mg of starting lysate) based on the Pierce 660 assay readings and protein abundances from shotgun proteomics analysis of trypsin digests of these samples by UT Southwestern’s (UTSW’s) proteomics core facility. Lysates (1 mg) were incubated with 250µL of magnetic streptavidin beads at 4°C for overnight incubation ∼16h. The next day, the beads were placed on a magnetic rack and flow throughs were saved. Beads were washed 2x with 200µL of 50mM Tris-HCl pH 7.5 followed by 2x with 2M urea + 50mM Tris-HCl pH 7.5. The beads were then incubated with 80µL 2M urea + 100μL 0.5ug/µl trypsin + 20µL 10mM DTT to achieve a final urea concentration of 1mM and a ratio of 1: 20 for trypsin: lysate, for 1h at 25°C with shaking at 1000rpm in a thermomixer. The beads were washed 2x with 60µl of 2M urea + 50mM Tris-HCl pH 7.5 and the two washes were combined with the supernatant. The eluate was reduced with DTT at a net concentration of 4mM by incubating for 30min with shaking at 1000rpm, 25°C. The samples were alkylated with 10mM iodoacetamide for 45min at 25°C with shaking at 1000rpm.

At the end of this reaction, 50mM Tris-HCl pH 7.5 was added to the solution to achieve a final urea concentration of 0.73M. The samples were incubated overnight (∼15h) at 37°C with shaking at 1000rpm to allow trypsin digestion to continue. The samples were removed from the thermomixer and spun down. Trypsin was quenched by acidifying the samples to pH <3 with the addition of formic acid at a final concentration of 1%.

### TMT Mass Spectrometry

5μL of 10% trifluoroacetic acid (TFA) was added to each sample, and solid-phase extraction was performed on each sample using an Oasis HLB 96-well uElution plate (Waters). The eluates were dried and reconstituted in 50μL of 100 mM triethylammonium bicarbonate (TEAB). 10μL of each sample was labeled with 4μL of a different TMT10plex reagent (Thermo Scientific, label TMT10-131 not used). Samples were quenched with 1μL of hydroxylamine, mixed, and dried in a SpeedVac. Samples were reconstituted in 2% acetonitrile, 0.1% formic acid to a concentration of 0.5 ug/μL,

2μL of each TMT sample were injected onto an Orbitrap Fusion Lumos mass spectrometer coupled to an Ultimate 3000 RSLC-Nano liquid chromatography system. Samples were injected onto a 75μm i.d., 75-cm long EasySpray column (Thermo) and eluted with a gradient from 0-28% buffer B over 180 min. Buffer A contained 2% (v/v) acetonitrile (ACN) and 0.1% formic acid in water, and buffer B contained 80% (v/v) ACN, 10% (v/v) trifluoroethanol, and 0.1% formic acid in water. The mass spectrometer was operated in positive ion mode with a source voltage of 1.8 kV and an ion transfer tube temperature of 275°C. MS scans were acquired at 120,000 resolution in the Orbitrap and top speed mode was used for SPS-MS3 analysis with a cycle time of 2.5 s. MS2 was performed with CID with a collision energy of 35%. The top 10 fragments were selected for MS3 fragmentation using HCD, with a collision energy of 55%. Dynamic exclusion was set for 25 s after an ion was selected for fragmentation.

### Proteomics Data Analysis

Raw MS data files were analyzed using Proteome Discoverer v2.4 (Thermo), with peptide identification performed using Sequest HT searching against the mouse protein database from UniProt. Fragment and precursor tolerances of 10 ppm and 0.6 Da were specified, and three missed cleavages were allowed. Carbamidomethylation of Cys and TMT labeling of N-termini and Lys sidechains were set as a fixed modification, with oxidation of Met set as a variable modification. The false-discovery rate (FDR) cutoff was 1% for all peptides.

For every biological replicate, absolute abundance of each protein was first normalized to the total protein abundance of a particular lysate sample to account for any differences in total protein concentrations across samples before comparison. These values were used to calculate the relative enrichment of proteins specific to tau seeding such as

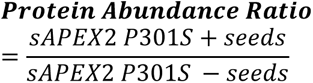

The relative values for sAPEX P301S cell line were compared to the relative values of the sAPEX only cell line (negative control) using the unpaired t-test, two-stage step-up ((Benjamini, Krieger, and Yekutieli), FDR 1.00%), on three independent biological replicates.

### Biosensor Seeding Assay

All biosensor assays were performed with naked seeding (no transfection reagent). Briefly, v2L cells and genetic modifications thereof were plated at a density of 15,000 cells/well of a 96 well plate and allowed to settle overnight. Cells were treated with appropriate concentration of recombinant tau fibrils for 48 hours upon which cells were harvested for flow cytometry. Fibril prep was sonicated in a water bath sonicator for 1min at 65Amp prior to cell treatment. Recombinant tau fibrils were prepared as previously described (31).

For seeding with brain lysates and for α-synuclein biosensor seeding assay, 8,000 cells/well were plated in 96 well plates and seeding was monitored for 72 hours. In the case of brain homogenates, biosensors were treated with 25μg of the lysate that was sonicated for 1min at 65 Amp in a water bath sonicator. For seeding with α-synuclein, fibrils were sonicated for 5min total, 1min on /1min off at 65 Amp, and used at a concentration of 400nM.

All seeding results have been reported as % FRET+ cells. Except for the cofactor data graphs, the FRET data has been plotted after subtracting the background signal (no exogenous tau added) which was negligible for all conditions (no FRET recorded in the absence of tau seeds) unless otherwise specified.

### Brain Homogenization

Brain tissue from clinically and neuropathologically well-characterized cases of AD and CBD were obtained from UTSW and Washington University in St. Louis. All human tissues used in these experiments were derived from autopsy subjects. Since deceased subjects are not considered human subjects for research purposes, these studies are considered exempt from human subjects research regulations and do not require IRB approval. Brain samples were weighed and added to 1X TBS supplemented with cOmplete Ultra (Roche) protease inhibitor to prepare a 10% w/v solution. The brains were homogenized using a probe homogenizer to obtain a slurry that was sonicated for 15min total, 1min on/ 30sec off. The sonicated samples were centrifuged at 4°C for 15min at 21,300g. Protein concentration of the supernatant was measured using Pierce 660 assay and was subsequently used for naked seeding.

### Uptake Assay

Uptake assay was performed as previously described (31). Briefly, v2L cells were plated overnight at a density of 8,000 cells/well of a 96 well plate. Cells were treated with 25nM of AF-647 labeled tau fibrils or AF-647 dye alone as a negative control. After 4h of incubation with the fibrils, cells were harvested with 0.25% trypsin for flow cytometry.

The labeled fibrils used in this assay were obtained by incubating recombinant tau fibrils (8μM, 200μL) with lyophilized AF-647 dye (25μg) for 1h at room temperature (RT) followed by quenching the reaction with 100mM glycine and subsequent dialysis in a 3500kDa dialysis cassette. This protocol was followed as previously described in detail (31).

The median fluorescence intensity (MFI) values representing the amount of tau internalized were plotted after subtracting the background MFI of the dye alone signal for all conditions. These MFI values were then normalized relative to the appropriate control condition of the respective experiment (DMSO, NTG, or Scr ctrl).

### Drug Treatments

96 well plates were coated with PDL and incubated at 37°C for 3h followed by washout with PBS. v2L cells were plated at a density of 15,000 cells/well and allowed to settle overnight. Cells were treated with different drugs (ML-240, NMS-873, and MG132) for about an hour upon which 25nM of recombinant tau fibrils were introduced to the media. After four hours of incubation with the fibrils (five hours with drugs), the media was replaced with fresh media and seeding or uptake was monitored for 48h and 4h, respectively.

### Flow Cytometry

To harvest cells for flow cytometry, media was removed, and cells were treated with 0.05% trypsin (0.25% trypsin for uptake assay) for 5min at 37°C (0.25% trypsin, 15min at 37°C in case of PDL coated plates). Trypsin was quenched with cold media and cells were resuspended a few times before transferring the suspension to 96-well round-bottom plates which were centrifuged at 1000 rpm for 5min. Supernatant was removed and the cell pellets were resuspended in 2% paraformaldehyde (PFA) and allowed to fix for 10min at RT. Cells were spun down again, PFA was removed, and cells were resuspended in PBS and stored at 4°C until ready to be run on a flow cytometer for quantifying the FRET signal.

### Cloning

FM5 vector with UBC promoter was used to clone all the APEX constructs. sAPEX fragments (AP and EX) were PCR amplified from the constructs provided by Dr. Alice Ting’s lab. Amplified sAPEX fragments were appended on to the c-terminus of RD tau fragments via a linker using overlap PCR. Using Gibson assembly, the final gene fragments were cloned into FM5 UBC plasmid which was cut with Esp3I. All Gibson reaction products were transformed into Stbl3 bacterial cells. Bacterial colonies were inoculated, DNA was purified using Qiagen miniprep kit, and the sequences were verified using Sanger sequencing at UTSW’s sequencing facility.

### Lentivirus Production

Low passage HEK293T cells were plated at ∼ 70% confluency in 6 well plates and allowed to settle overnight. A master mix was prepared using 400ng of plasmid of interest, 400ng of VSVG, and 1200ng of PSP plasmids required for virus packaging, along with 7.5μL of TransIT 293T and 120ul of OMEM per well of a 6 well plate. The master mix was allowed to incubate at RT for 30min upon which it was added to the cells in a drop-wise fashion. The virus was harvested 48h later by collecting the media, spinning it for 5min at 1000rpm, and then freezing the aliquoted supernatant.

### CRISPR/Cas9 screen for VCP cofactors

CRISPR constructs for the cofactors were outsourced to Twist Biosciences for synthesis. Constructs not synthesized by the company were cloned in the lab using standard ligation reaction. Four guides per gene were chosen from the Brunello library deposited online and ordered as duplex DNA from IDT. LentiCRISPRv2 (Addgene #52961) was cut using Esp3I, and T4 ligase was used for all ligation reactions of the guides into the plasmid. Stbl3 bacteria were transformed with the ligated products, selected colonies were inoculated, mini-prepped using Qiagen kit, and the purified DNA was sequence-verified. Pooled lentivirus was prepared with four constructs per gene and v2L cells were transduced with the virus at the desired MOI. After 24h, cells were expanded in puromycin media (2μg/mL) for selection of the KO population. Selected populations were eventually used for seeding and uptake assays and western blot to ensure the gene was knocked out.

### siRNA knockdown

siRNAs were ordered from Origene. 300,000 v2L cells were plated in 6 well plates and allowed to settle overnight. The next day, cells were treated with 100nM of each siRNA, with a total of three siRNAs per gene using RNAiMax Lipofectamine (Thermo) as a transfection vehicle at 7.5µl/well. After 48h of transfection, the cells were plated in 96 well plates for seeding and uptake assays. Cells were also used for western blot to verify the knockdown.

### Western blot

Cell pellets were lysed in RIPA buffer and allowed to sit on ice for 5min followed by a 15,000g spin for 10min at 4°C. The supernatants were used to determine the protein concentrations using Pierce 660 assay. 15μg of total protein was treated with SDS buffer and heated at 95°C for 10min. Samples were loaded onto 4-12% bis-tris gels and the proteins were transferred onto nitrocellulose membranes using the Biorad turbo transfer machine. All incubations for subsequent steps were done in TBS + 0.05% Tween-20 (TBST). The membranes were first incubated in blocking buffer (5% milk powder + TBST) for 1h at RT, followed by primary antibodies in the blocking buffer at 4°C with overnight shaking. After the primary antibody incubation, the membranes were washed 3x with TBST, 10min each. Then, appropriate HRP-conjugated secondary antibodies in blocking buffer were added to the membranes for a 1.5h incubation at RT. Membranes were again washed 3x in TBST followed by a single wash in TBS alone before reading the HRP signal using the Thermo ECL kit.

### Statistical Analysis

Statistical analyses were performed using GraphPad Prism. One-Way ANOVA (Šídák method) with a 95% confidence interval was used for all statistical analyses unless otherwise stated. The P values are described as follows: ns = not significant/ P>0.05, * = P ≤ 0.05, ** = P ≤ 0.01, *** = P ≤ 0.001, **** = P ≤ 0.0001.

### Graphics

Biorender.com was used to create the graphics presented here.

## Declarations

### Ethics approval and consent to participate

Not applicable.

### Consent for publication

The authors give consent for publication.

### Availability of Data

Data generated in this study and not presented here are available from the corresponding author on request.

### Author contributions in this study

S.B, J.V.A, and M.I.D designed the study. S.B did the experiments. J.V.A and V.A.M assisted with cloning. O.K prepped the tau fibrils. A.L and his core facility assisted with mass spectrometry. N.J.C and C.L.W provided samples. S.B, J.V.A and M.I.D wrote the manuscript.

## Acknowledgements

We are grateful to Dr. Alice Ting for providing us with the sAPEX2 constructs prior to their own publication, allowing us a head start on our project. We are also thankful to Dr. Chris Weihl for sharing his VCP insights with us as we ventured into a new area of research. Drs. Donna Huryn, Peter Wipf, and Ray Deshaies advised on our results related to VCP inhibitors. Special thanks to Dr. Sandra Schmid for her critical discussions of the findings of this study.

## List of Reagents

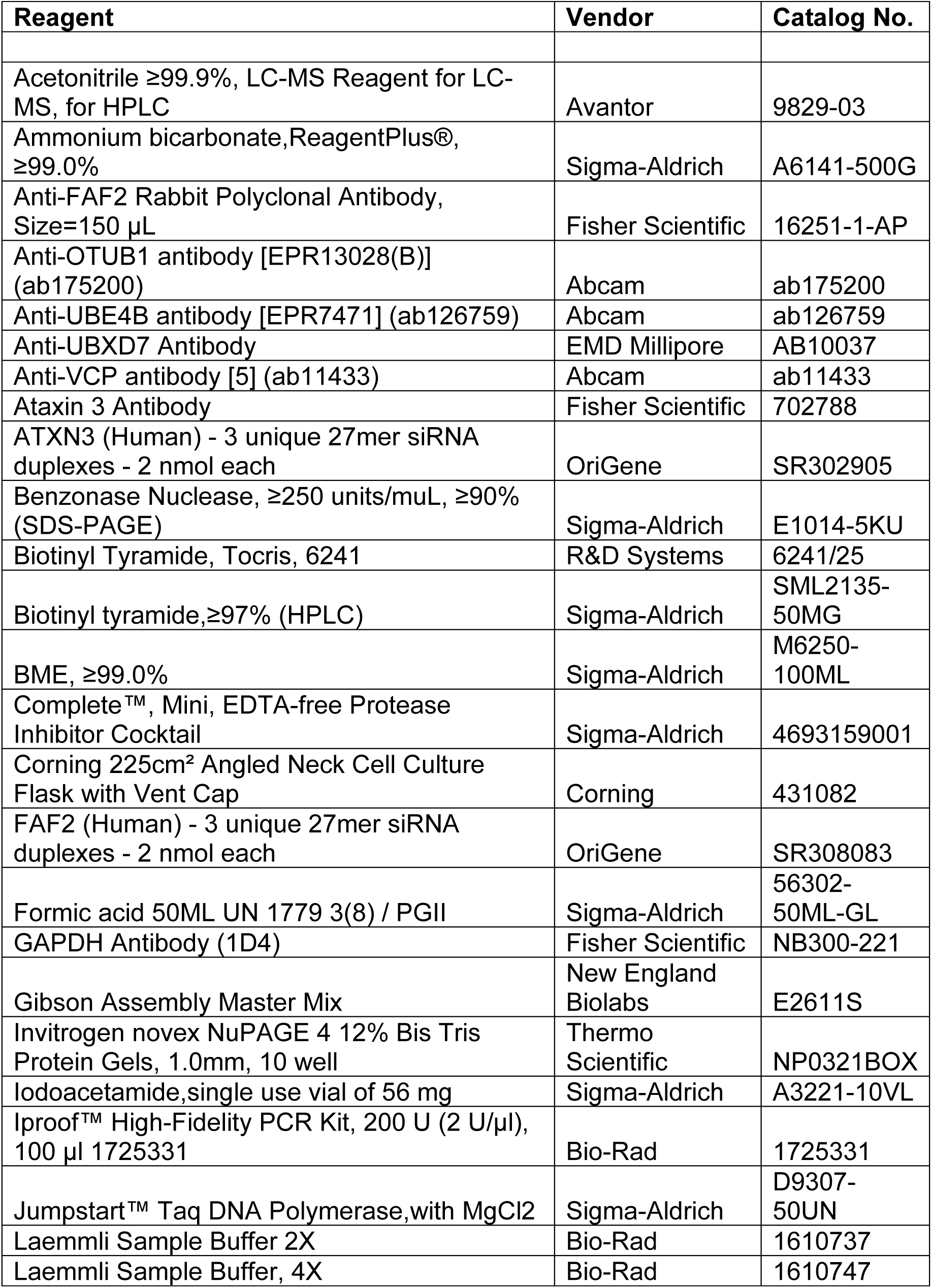
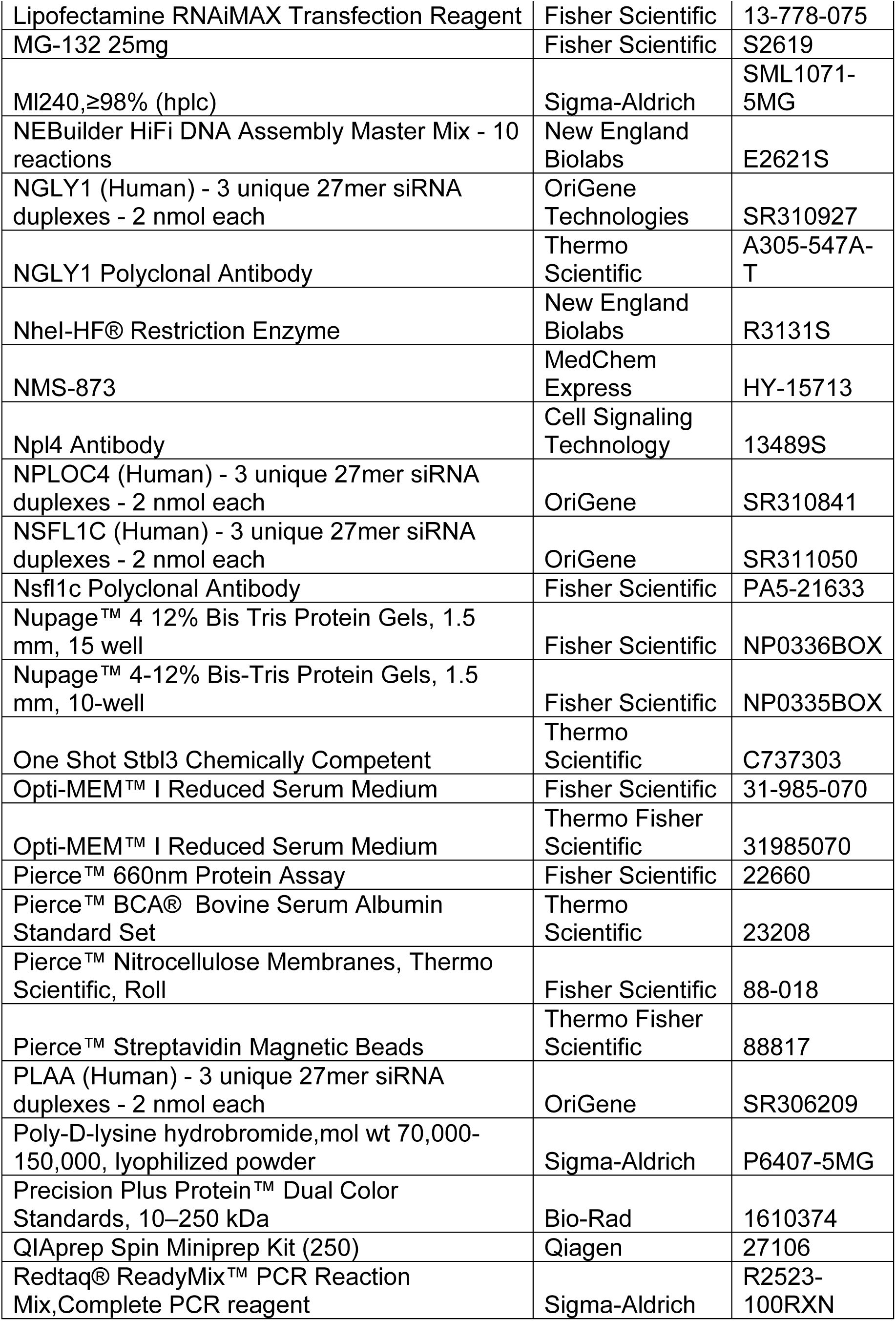
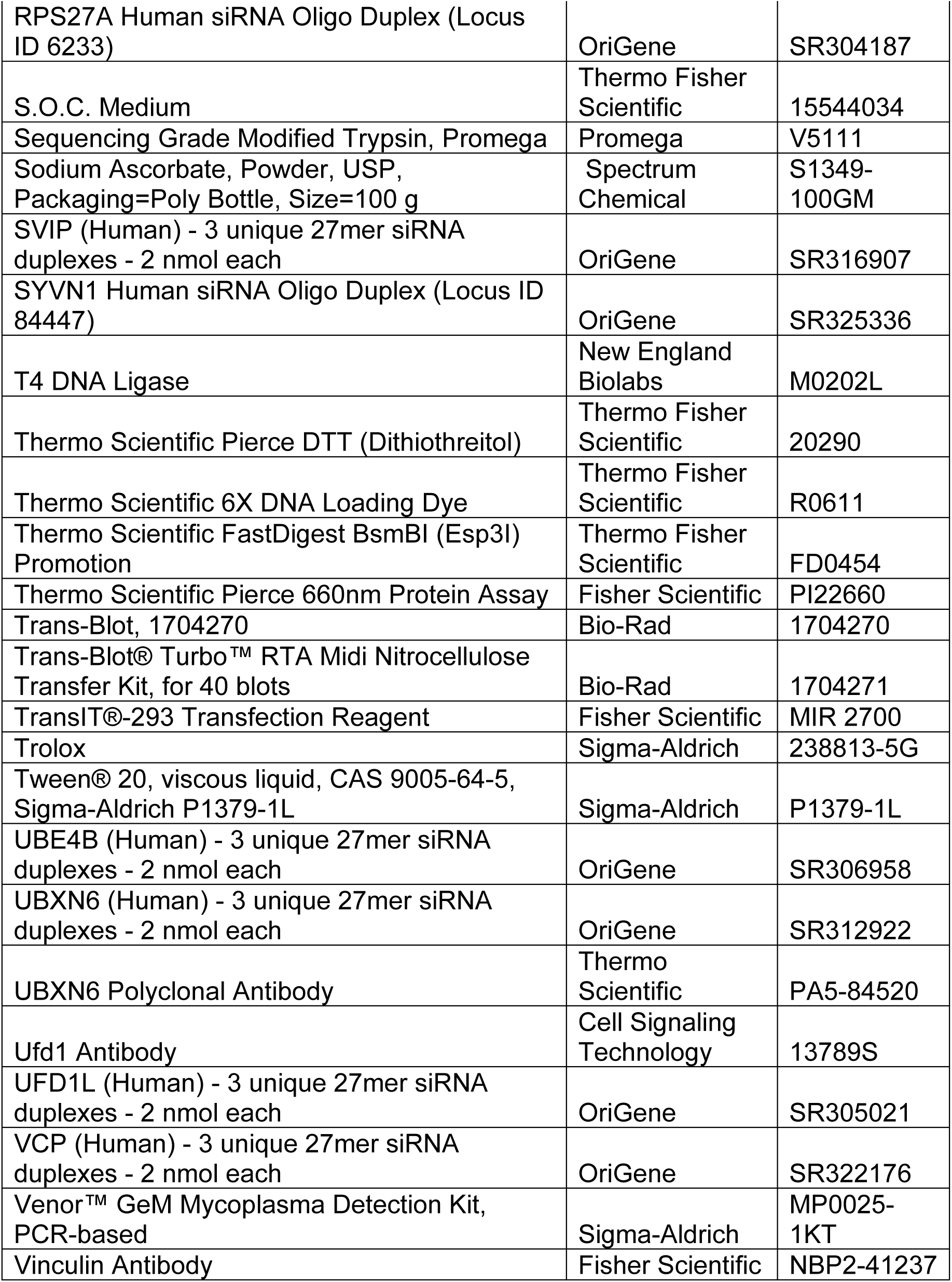

## List of abbreviations

VCP/p97: valosin containing protein
sAPEX2: split ascorbate peroxidase
2 RD: repeat domain
FL: full length
WT: wild type
BP: biotin-phenol
TMT-MS: tandem mass-tag mass spectrometry
KO: knockout
KD: knockdown
AF-647: Alexa fluor-647
AD: Alzheimer’s disease
CBD: corticobasal degeneration

---

**Supplemental Figure 1.**
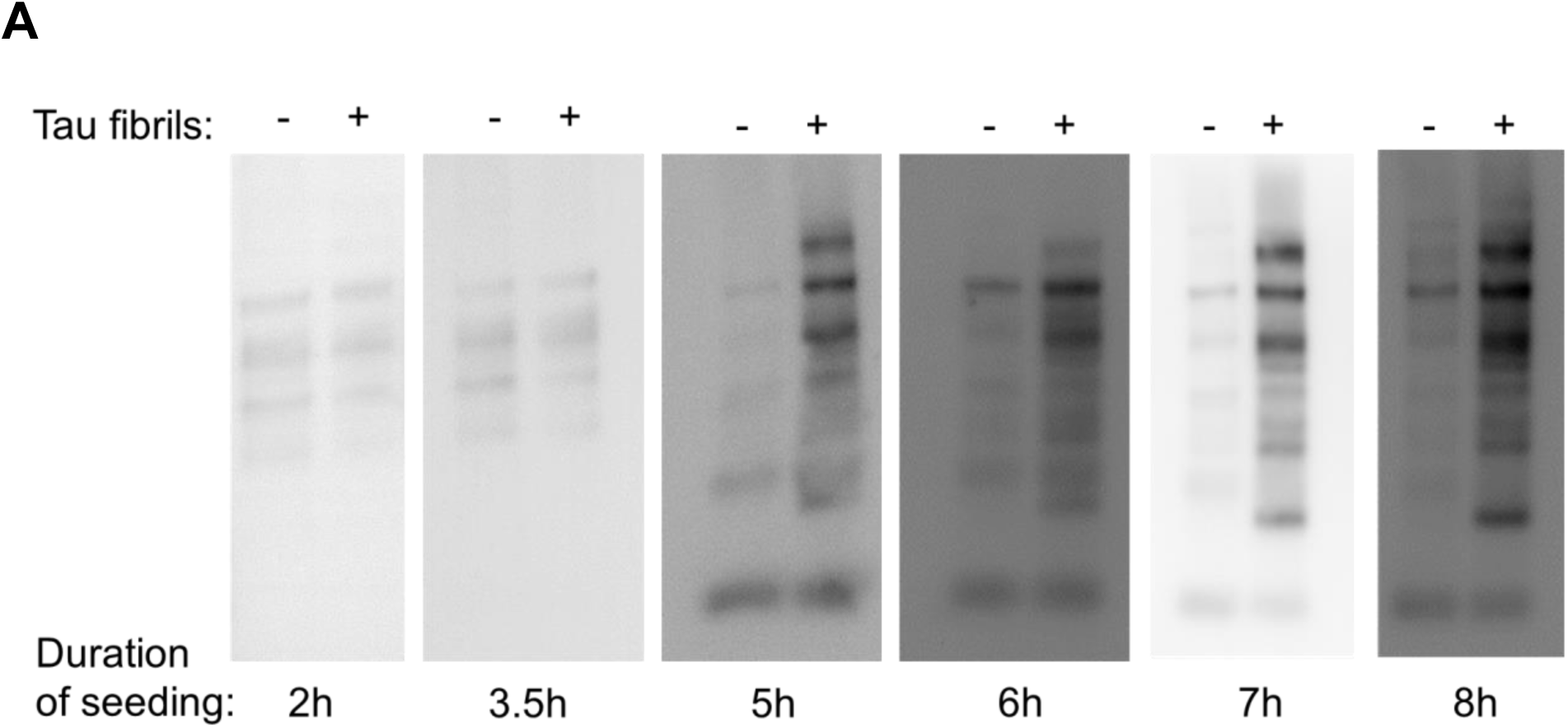
Proximity labeling of nascent tau aggregates identifies VCP. **(A)** Western blot probed for biotinylation signal using streptavidin-HRP shows the earliest reconstitution of P301S tau-sAPEX2 activity at 5h.

**Supplemental Figure 2.**
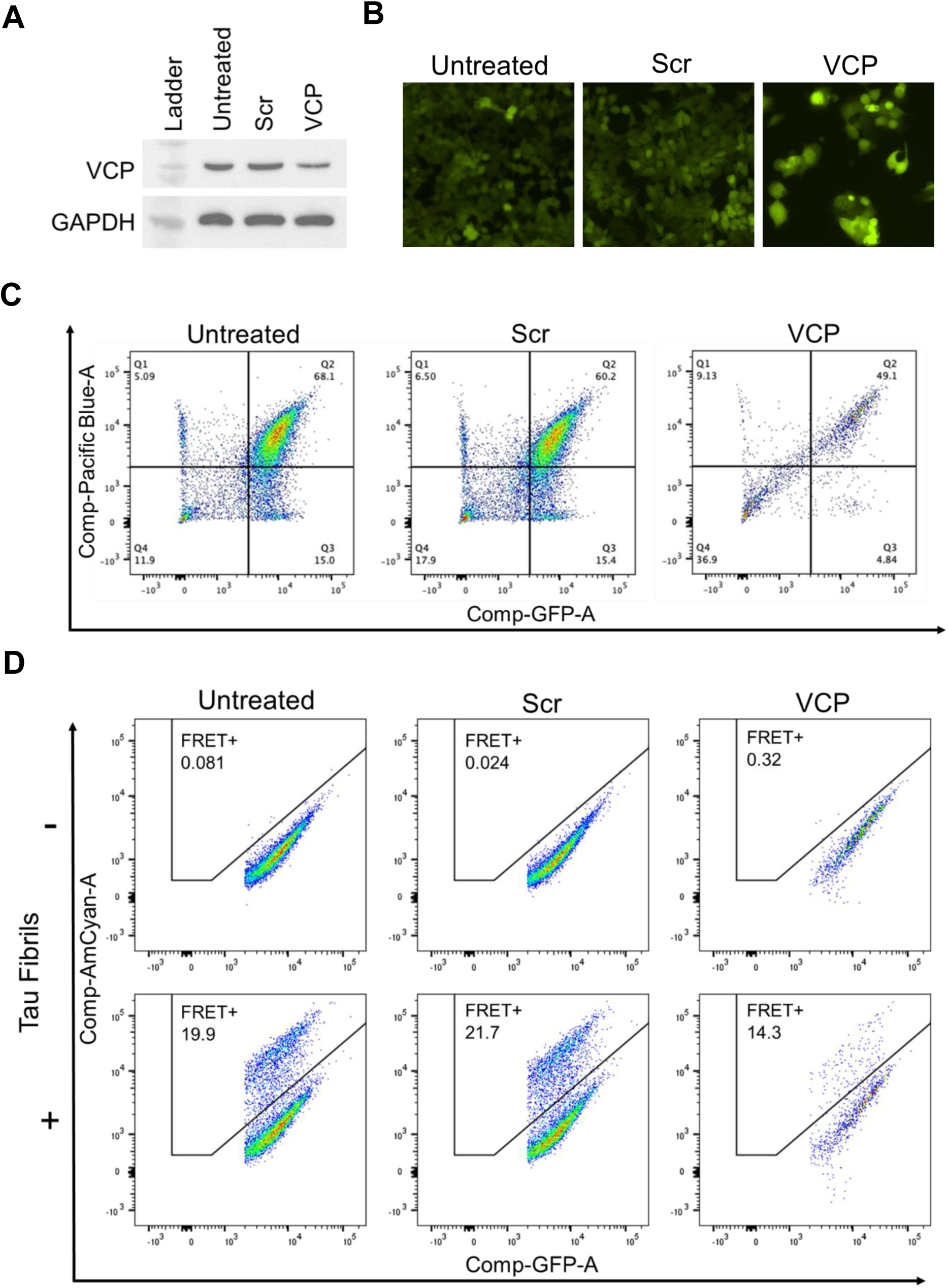
Genetic reduction of VCP reduces tau seeding. **(A)** Western blot showing KD of VCP compared to scrambled (Scr) control siRNA treated cells. **(B)** Images representing tau-clover signal as observed by fluorescence microscopy (20x). VCP KD cells were brighter but fewer in number due to reduced cell proliferation. **(C)** Flow plots depicting a shift in dual positive biosensor population in quadrant 2 (Q2) for the VCP KD cell line highlighted the increase in fluorescence levels of the biosensors as observed under the microscope. Cell proliferation was reduced. **(D)** Flow plots showing no background spontaneous seeding (FRET+ values) in the VCP KD cells in the absence of exogenous tau fibrils, despite the increased basal fluorescence.

**Supplemental Figure 3.**
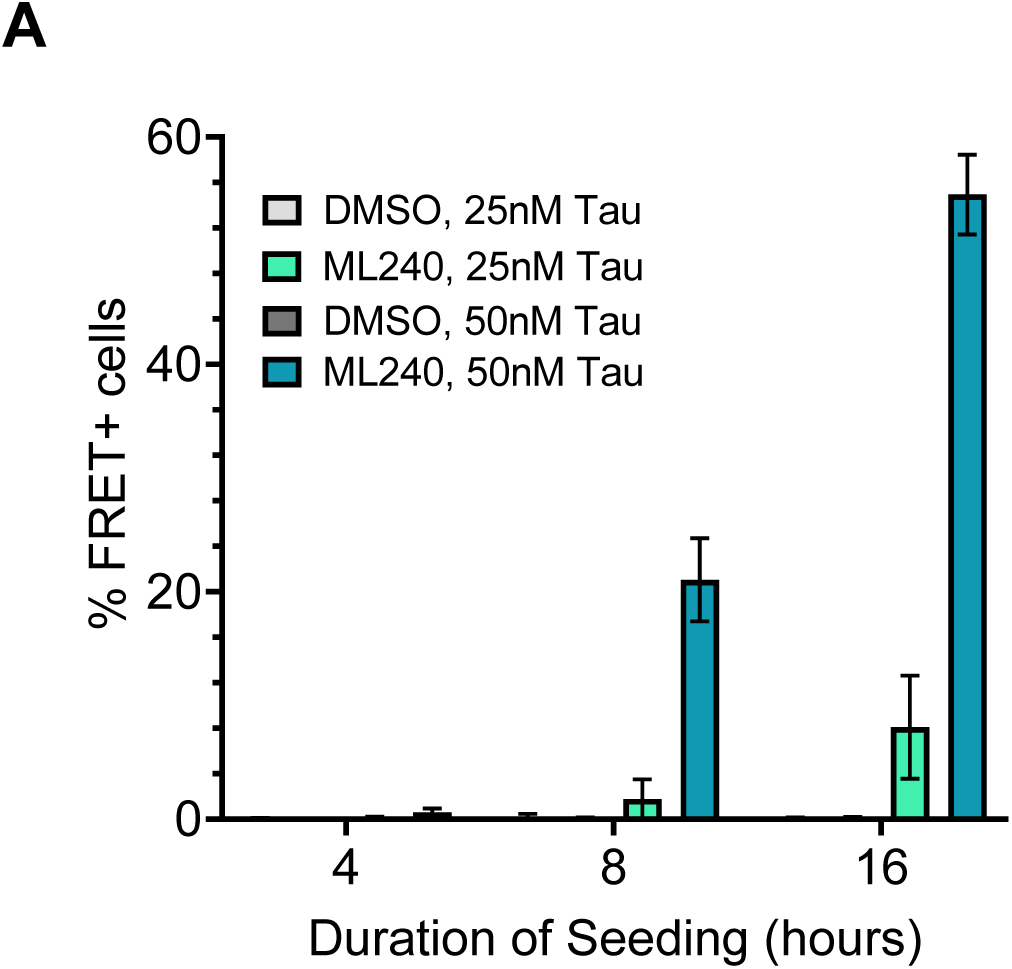
Acute exposure of inhibitors differentially impacts tau aggregation. **(A)** ML-240 increased the kinetics of tau seeding with a robust FRET signal detectable as early as 8h.

**Supplemental Figure 4.**
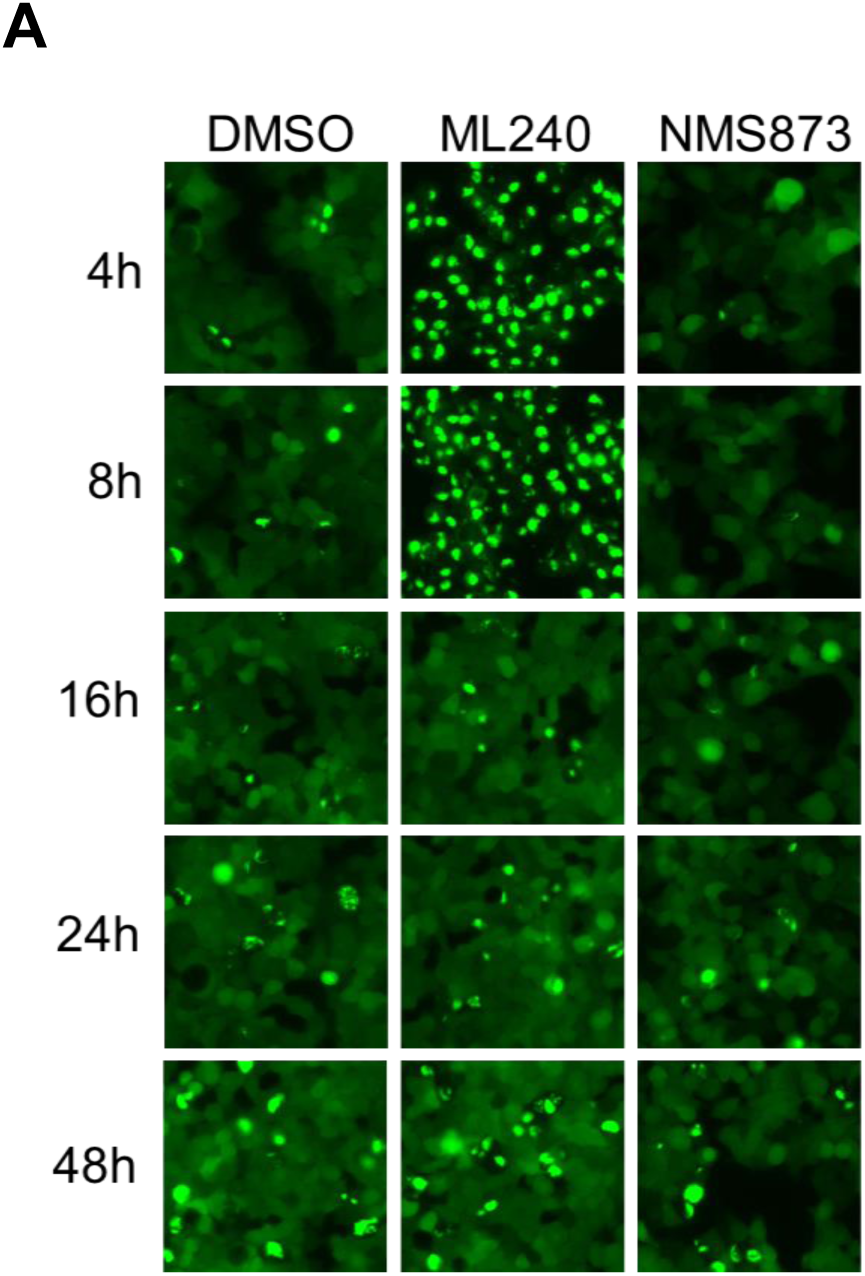
Only early VCP inhibition impacts tau seeding. **(A)** ML-240 increased, and NMS-873 decreased tau aggregation only within the initial 8h of seeding as represented by tau-clover images (20x magnification) at different time points of the seeding process.

**Supplemental Figure 5.**
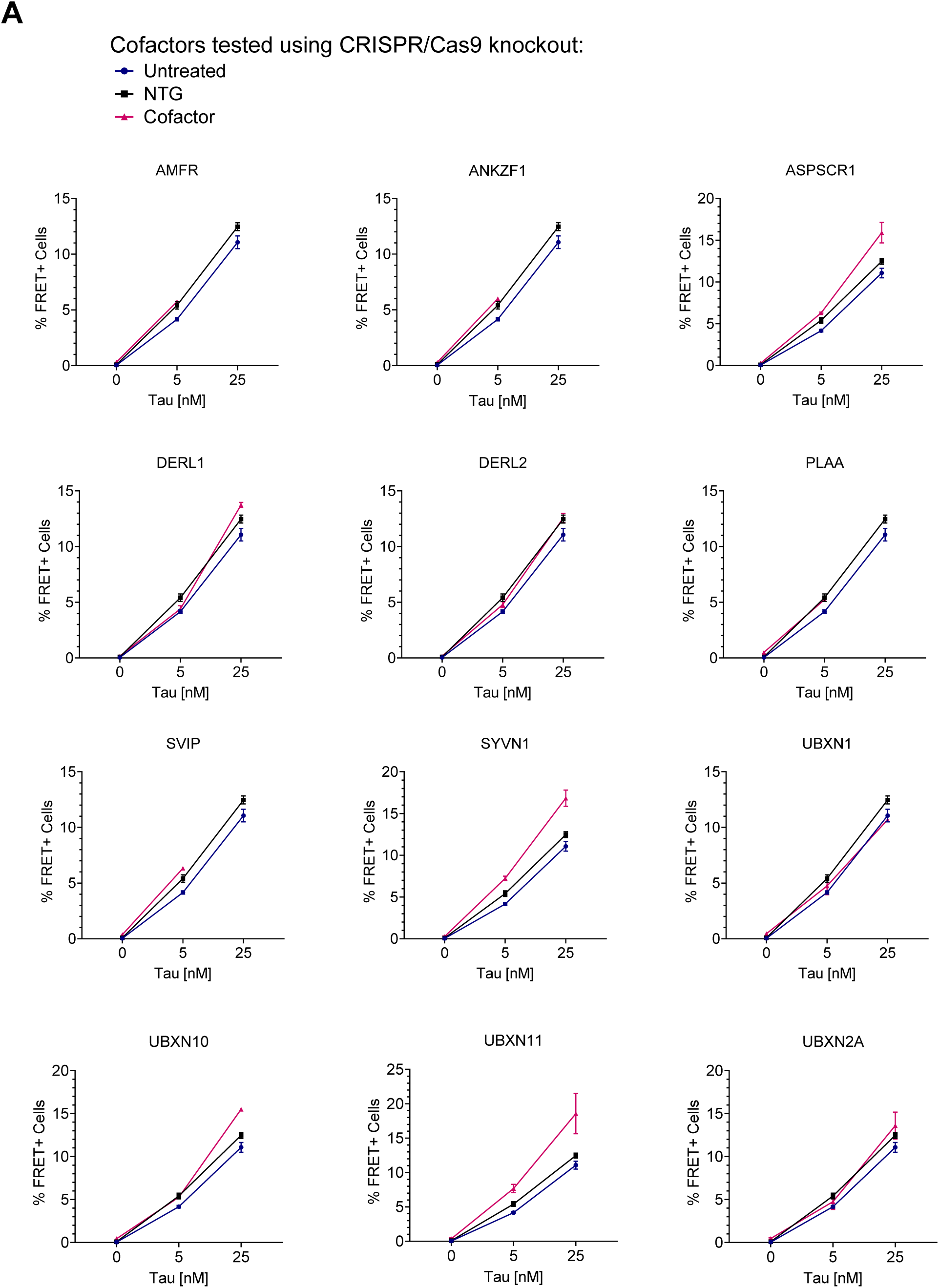
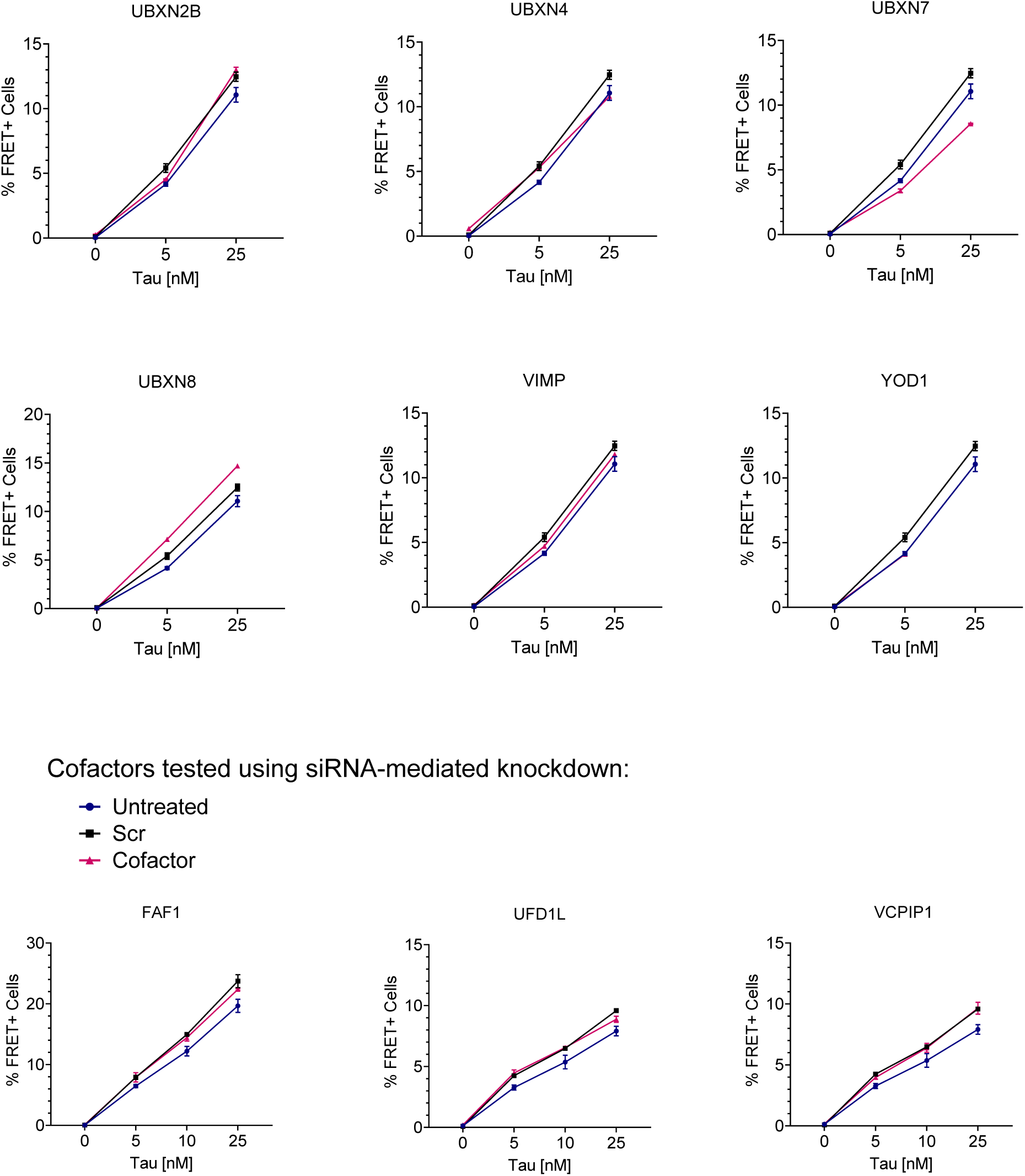
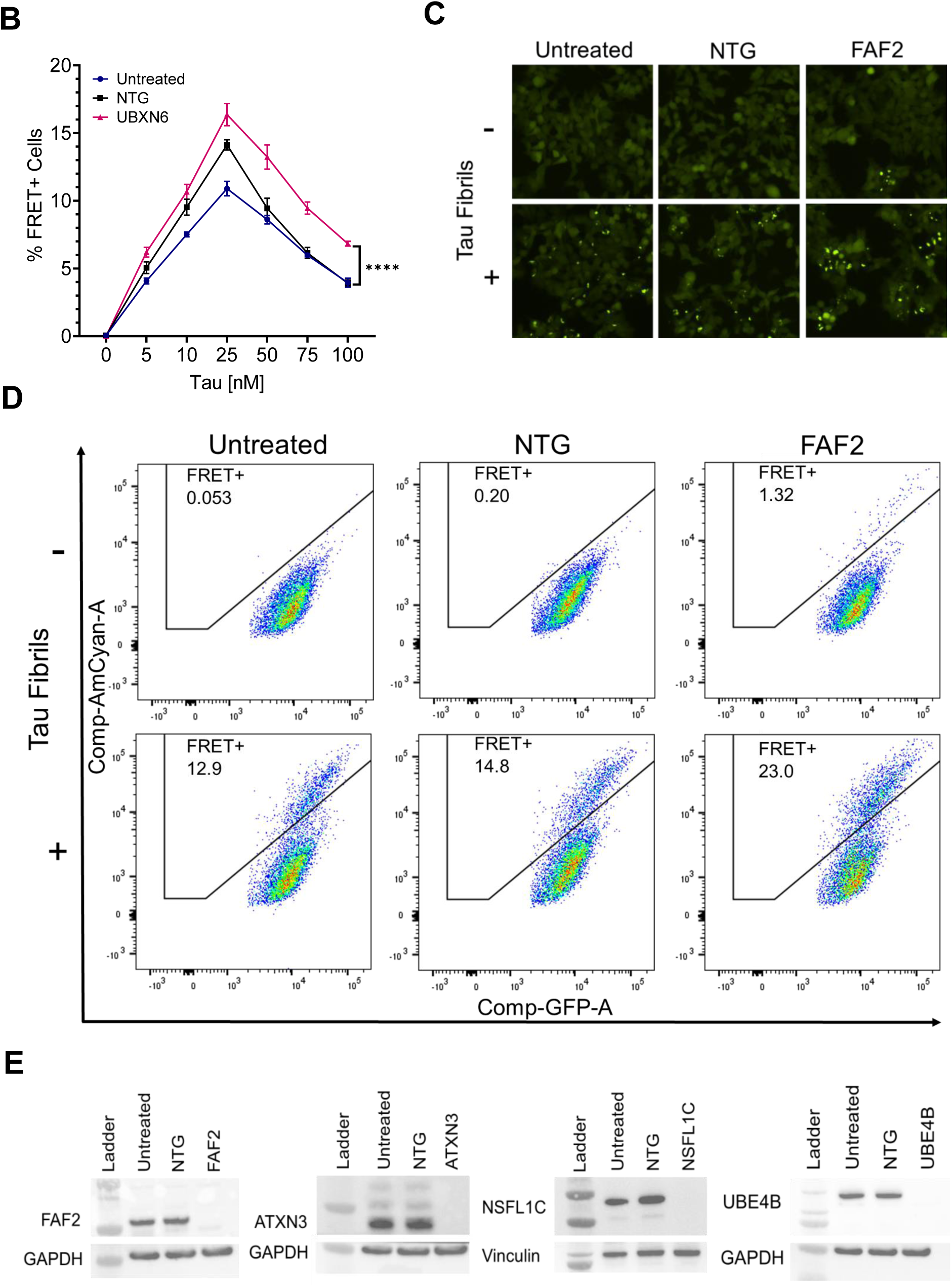
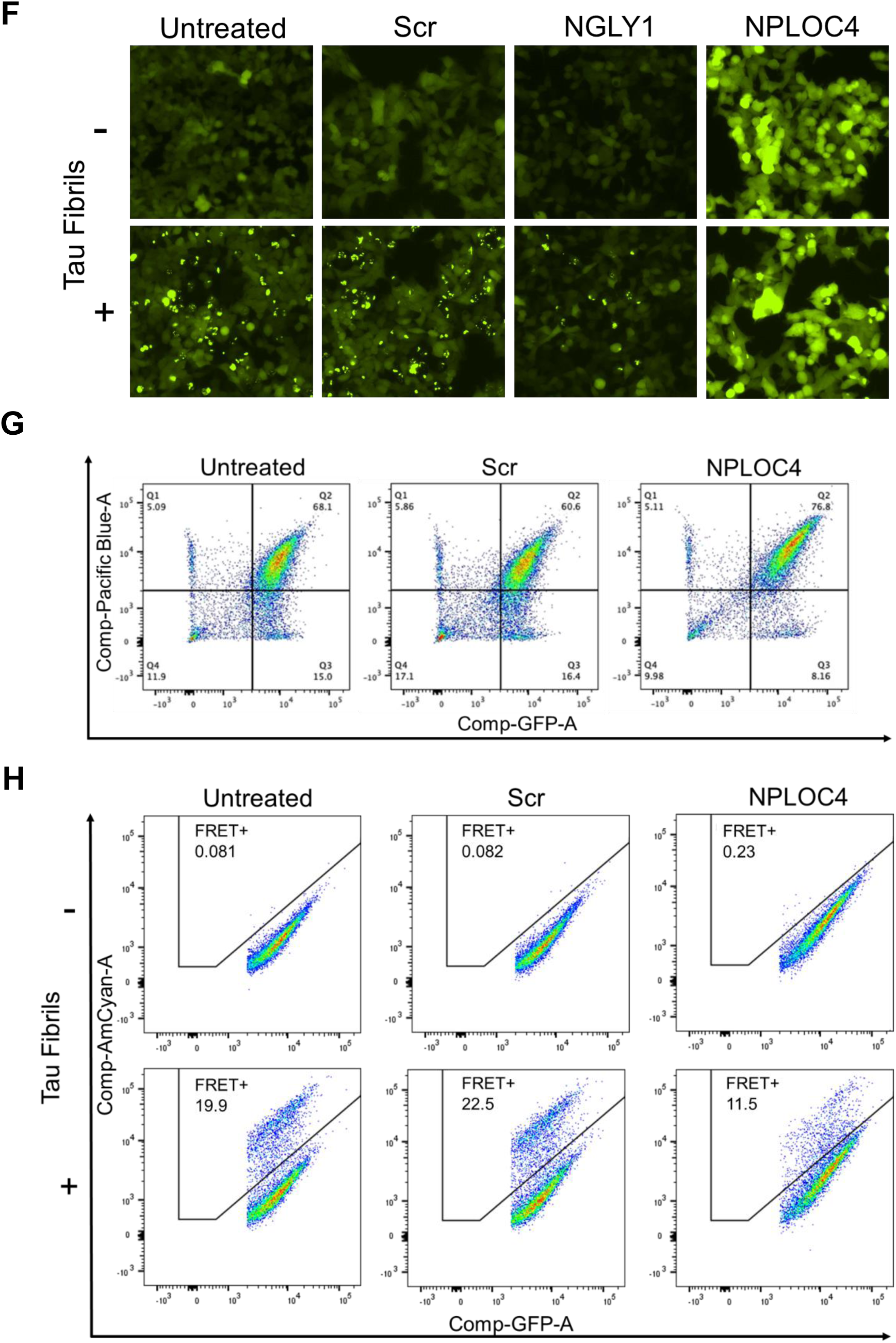
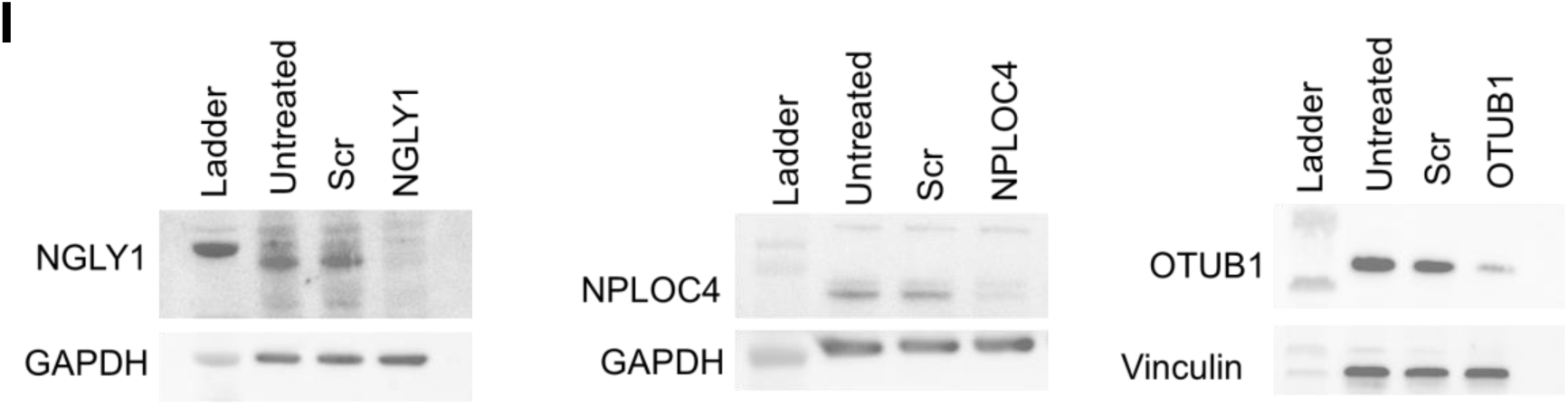
VCP cofactors differentially regulate tau aggregation. **(A)** Graphs representing the % FRET+ signal for cofactors (KO and KD) that did not affect tau seeding. **(B)** KO of UBXN6 increased tau seeding with an effect most pronounced at higher tau concentrations. Error bars represent S.D. Graph representative of n=3 independent experiments. One-Way ANOVA (Šídák method) with a 95% confidence interval, P Value **** < 0.0001. **(C)** KO of FAF2 caused spontaneous aggregation as observed by tau-clover puncta (20x magnification). **(D)** FRET flow cytometry documented spontaneous aggregation in the FAF2 KO cells in the absence of exogenously added tau seeds. **(E)** Western blots showing absence of FAF2, ATXN3, NSFL1C, and UBE4B in the respective knockout cells lines. Non-targeting guide (NTG) treated cell line was used as a negative control. **(F)** Representative images showing increased basal fluorescence in NPLOC4 KD biosensors. **(G)** Flow cytometry indicated a shift in dual positive biosensor population in quadrant 2 (Q2) for the NPLOC4 KD cell line highlighting the increase in fluorescence levels of the biosensors as also observed under the microscope. **(H)** FRET flow cytometry revealed no spontaneous background aggregation in the NPLOC4 KD cells in the absence of exogenous tau fibrils, despite increased basal fluorescence. **(I)** Western blots showing reduced protein levels of the cofactors in their respective KD cell lines. Scrambled siRNA (Scr) treated cell line was a negative control.

